# Thousands of novel unannotated proteins expand the MHC I immunopeptidome in cancer

**DOI:** 10.1101/2020.02.12.945840

**Authors:** Tamara Ouspenskaia, Travis Law, Karl R. Clauser, Susan Klaeger, Siranush Sarkizova, François Aguet, Bo Li, Elena Christian, Binyamin A. Knisbacher, Phuong M. Le, Christina R. Hartigan, Hasmik Keshishian, Annie Apffel, Giacomo Oliveira, Wandi Zhang, Yuen Ting Chow, Zhe Ji, Irwin Jungreis, Sachet A. Shukla, Pavan Bachireddy, Manolis Kellis, Gad Getz, Nir Hacohen, Derin B. Keskin, Steven A. Carr, Catherine J. Wu, Aviv Regev

## Abstract

Tumor epitopes – peptides that are presented on surface-bound MHC I proteins - provide targets for cancer immunotherapy and have been identified extensively in the annotated protein-coding regions of the genome. Motivated by the recent discovery of translated novel unannotated open reading frames (nuORFs) using ribosome profiling (Ribo-seq), we hypothesized that cancer-associated processes could generate nuORFs that can serve as a new source of tumor antigens that harbor somatic mutations or show tumor-specific expression. To identify cancer-specific nuORFs, we generated Ribo-seq profiles for 29 malignant and healthy samples, developed a sensitive analytic approach for hierarchical ORF prediction, and constructed a high-confidence database of translated nuORFs across tissues. Peptides from 3,555 unique translated nuORFs were presented on MHC I, based on analysis of an extensive dataset of MHC I-bound peptides detected by mass spectrometry, with >20-fold more nuORF peptides detected in the MHC I immunopeptidomes compared to whole proteomes. We further detected somatic mutations in nuORFs of cancer samples and identified nuORFs with tumor-specific translation in melanoma, chronic lymphocytic leukemia and glioblastoma. NuORFs thus expand the pool of MHC I-presented, tumor-specific peptides, targetable by immunotherapies.

## Introduction

The major histocompatibility complex class I (MHC I) immunopeptidome consists of thousands of short 8-12 amino acid peptide antigens displayed on the cell surface by MHC I molecules. “Non-self” antigens presented by MHC molecules are recognized by CD8 T cells that mount an immune response. This defense mechanism can be exploited to target the immune system against cancer cells, which display cancer-specific antigens (neoantigens) on MHC I (Hu, Ott, and Wu 2018). Patients immunized with neoantigen-based vaccines display expanded neoantigen-specific T cells, suggesting that this could be a promising therapeutic avenue (Hilf et al. 2019; Keskin et al. 2019; Ott et al. 2017; Sahin et al. 2017).

Neoantigens are currently predicted based on the detection of cancer-specific somatic mutations in annotated protein-coding regions by whole exome sequencing (WES) and RNA-seq (Hilf et al. 2019; Keskin et al. 2019; Ott et al. 2017; Sahin et al. 2017). This approach often falls short for patients with few somatic mutations, generating few actionable neoantigens(Rajasagi et al. 2014). Several lines of evidence suggest that the potential sources of neoantigens are more varied. First, immune responses have been detected against antigens derived from retained introns, alternative open reading frames (ORFs) within coding genes and antisense transcripts (Robbins et al. 1997; Van Den Eynde et al. 1999; Wang et al. 1998). Additionally, while the MHC I immunopeptidome mainly consists of peptides derived from homeostatic protein turnover (Abelin et al. 2017; Sarkizova et al. 2019), peptides can also be sourced from alternative precursors, such as defective ribosomal products (DRiPs), and presumably “non-coding” regions of the genome (Laumont et al. 2016, 2018; Yewdell 2011). In particular, ribosome profiling (Ribo-seq), which assays mRNA translation by capturing and sequencing ribosome-protected mRNA fragments (RPFs) (Ingolia et al. 2009), has detected a plethora of translated novel unannotated open reading frames (nuORFs).

These nuORFs are derived from the 5’ and 3’ untranslated regions (UTRs), overlapping yet out-of-frame alternative ORFs in annotated protein-coding genes, long non-coding RNAs (lncRNAs), pseudogenes and other transcripts currently annotated as non-protein coding (Fields et al. 2015; Ji et al. 2015; Chew et al. 2013). Ribo-seq analysis of HEK293T, HeLa-S3, and K562 cell lines and of human fibroblasts infected with HSV-1 and HCMV has identified translated nuORFs that contribute peptides to the MHC I immunopeptidome, suggesting that nuORFs could also have an immunological function (Erhard et al. 2018; Martinez et al. 2019). However, a global understanding of the extent to which nuORFs contribute to the immunopeptidomes of healthy and cancer tissues, as well as the diversity and tissue specificity of nuORFs is still lacking.

## Results

### A comprehensive pipeline for Ribo-seq based nuORF identification

We hypothesized that cancer-associated processes could lead to nuORFs that are either mutated or exhibit tumor-specific expression and thus could serve as sources of neoantigens. To systematically evaluate the contribution of nuORFs to the MHC I immunopeptidome, we identified translated nuORFs using Ribo-seq; built an ORF database appending nuORFs detected by Ribo-seq to known annotations; and used this updated database to search for presented nuORFs in MHC I immunopeptidome mass spectrometry (MS) data (**Figure 1a**). Because MS/MS spectra are traditionally searched against a sequence database of annotated proteins, any presented peptides derived exclusively from nuORFs (which are, by definition, not in the standard annotated protein database) will not be identified by conventional search strategies. Therefore, we reasoned that including Ribo-seq-detected nuORFs in the search space can improve MHC I immunopeptidome identification.

**Figure 1.**
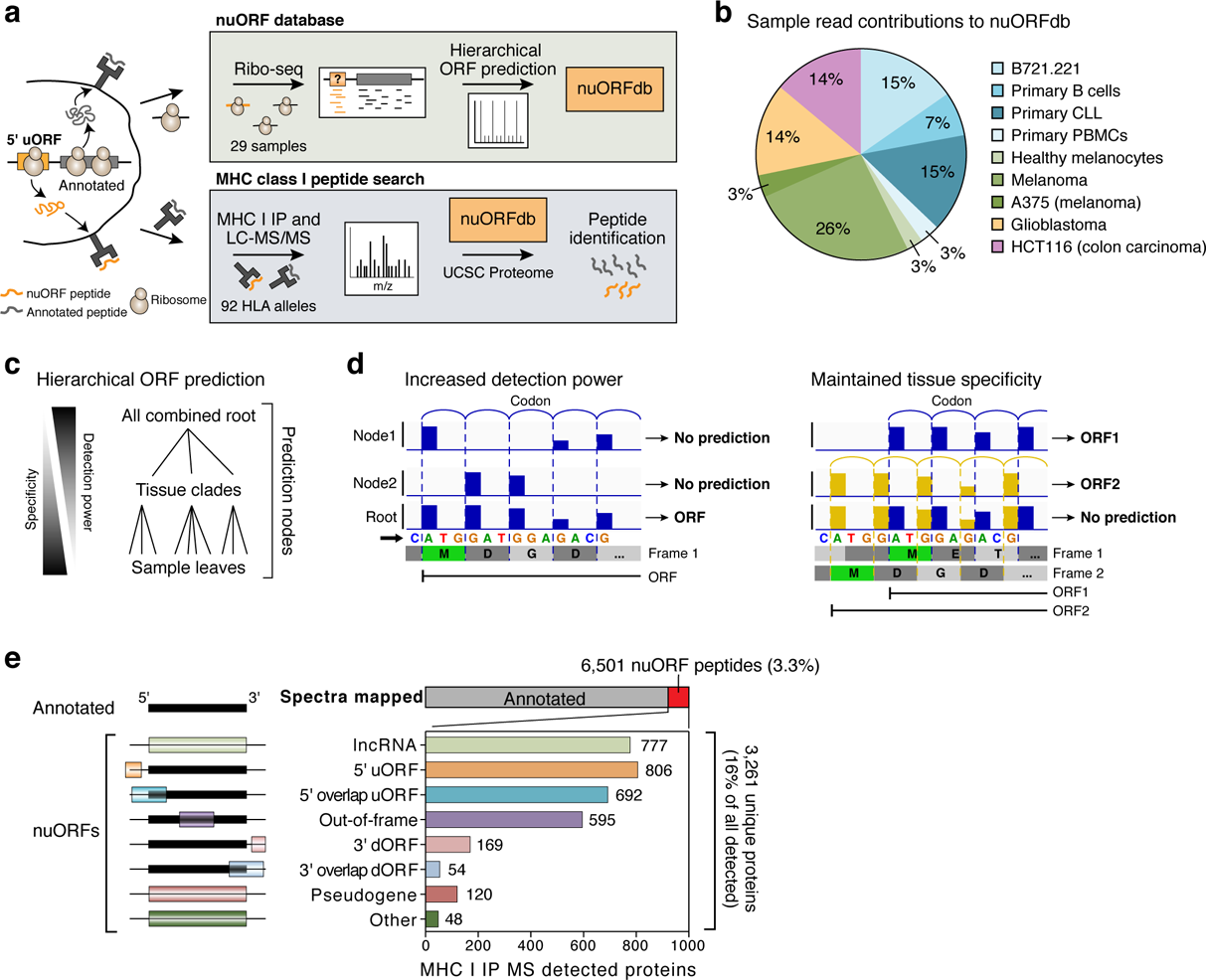
Thousands of nuORFs from Ribo-seq are translated and contribute peptides to the MHC I immunopeptidome. **a.** Schematic overview of nuORF database generation using Ribo-seq and hierarchical ORF prediction followed by nuORF peptide identification in MHC I immunopeptidomes. **b.** Sample read contribution to nuORFdb shown as percent of Ribo-seq reads contributed by each tissue type. **c.** Hierarchical ORF prediction approach. ORFs are predicted independently at multiple nodes from reads in each sample (leaves), multiple samples of the same tissue (clades) and all samples (root). **d.** Hierarchical prediction increases power while maintaining tissue specificity. Left: Pooling reads across samples allows ORF detection (bottom track) even when each sample alone will have insufficient reads (top two tracks). Right: Predicting in individual samples (top two tracks) detects overlapping ORFs. **e.** Diverse nuORFs contribute to the MHC I immunopeptidome. Top: Percent of MS/MS spectra mapped to nuORF peptides (red) identified in the MHC I immunopeptidome of 92 HLA mono-allelic B721.221 samples. Bottom: The number of detected nuORFs (x axis) of various types (y axis).

To this end, we first collected Ribo-seq data from 29 primary healthy and cancer samples and cell lines. These included primary normal and chronic lymphocytic leukemia (CLL) B cells, patient-derived primary glioblastoma (GBM) and melanoma cell cultures, primary healthy melanocytes, as well as established colon carcinoma and melanoma cell lines. These also included B721.221 cells, the parental cell line previously used to generate 92 single HLA allele-expressing lines from which we collected mono-allelic MHC I immunopeptidome data (Abelin et al. 2017; Sarkizova et al. 2019) (**Figure 1b**, **Supplementary Figure 1a**). We developed a hierarchical ORF prediction pipeline, where ORF predictions were carried out at multiple prediction nodes, consisting of each sample (leaf), tissue (clade) and across all samples combined (root) (**Figure 1c**, **Supplementary Figure 1a**, **Methods**). This approach aggregated signal across our Ribo-seq dataset to predict lowly translated ORFs, while maintaining sensitivity for tissue-specific, overlapping ORFs (**Figure 1d**). We predicted translated ORFs within transcripts annotated in GENCODE (Frankish et al. 2019) and in MiTranscriptome, which contains *de novo* assembled transcripts from thousands of RNA-seq libraries from tumors, normal tissues and cell lines, and therefore might be particularly useful to identify cancer-specific nuORFs (Iyer et al. 2015). Thus, we generated nuORFdb v1.0, containing 86,421 annotated and 237,427 nuORFs (323,848 ORFs in total). NuORFdb has ∼50-fold fewer ORFs than the ∼17 million ORFs obtained from the combined transcripts in the GENCODE and MiTranscriptome annotations (**Supplementary Figure 1b**). Compared to the annotated proteome (UCSC), nuORFdb has 1.46-fold more candidate MHC I-compatible 9mer peptides, making it practical for routine use in immunopeptidomics studies (**Supplementary Figure 1c**).

### NuORF derived peptides are presented on MHC I

Next, we searched the MHC I immunopeptidome MS/MS spectra from 92 HLA alleles expressed in B721.221 cells (Sarkizova et al. 2019) against nuORFdb with stringent FDR filtering (**Supplementary Figure 2**), and identified 6,501 high confidence (FDR<1%) peptides from 3,261 nuORFs, across various nuORF types (**Figure 1e**, **Supplementary Figure 3a**, **Methods**). NuORFs contributed 3.3% of peptides to the MHC I immunopeptidome, and 16% of all detected proteins with at least one MHC-presented peptide (**Figure 1e**).

Several lines of evidence revealed the MS/MS-identified nuORF peptides to be of comparable quality and characteristics to peptides from annotated ORFs (“annotated peptides”). First, nuORF and annotated MS/MS-detected peptides had similar Spectrum Mill MS/MS identification scores (11.7 nuORF *vs*. 11.4 annotated mean scores, 95% CI: 0.27-0.43), median peptide length (9AA), and translation levels (1.7 nuORF *vs*. 1.6 annotated mean log_2_TPM, 95% CI: 0.09-0.19) (**Figure 2a-c**, **Supplementary Figure 3b-d**). Second, chromatographic retention times for nuORF peptides correlated as well with predicted hydrophobicity indices as they did for annotated peptides (p=0.55, rank-sum test) (**Figure 2d**, **Supplementary Figure 3e**) (Mylonas et al. 2018; Rolfs et al. 2019). Finally, two-dimensional projection of pairwise distances amongst detected peptide sequences per allele showed that anchor residue motifs of nuORF-derived peptides matched closely to peptides derived from annotated proteins (**Figure 2e**, **Supplementary Figure 3f,g**) with a strong agreement in peptide sequences across all alleles (Pearson *r*^2^ = 0.85 nuORFs, *r*^2^ = 0.92 annotated) (**Figure 2f**).

**Figure 2.**
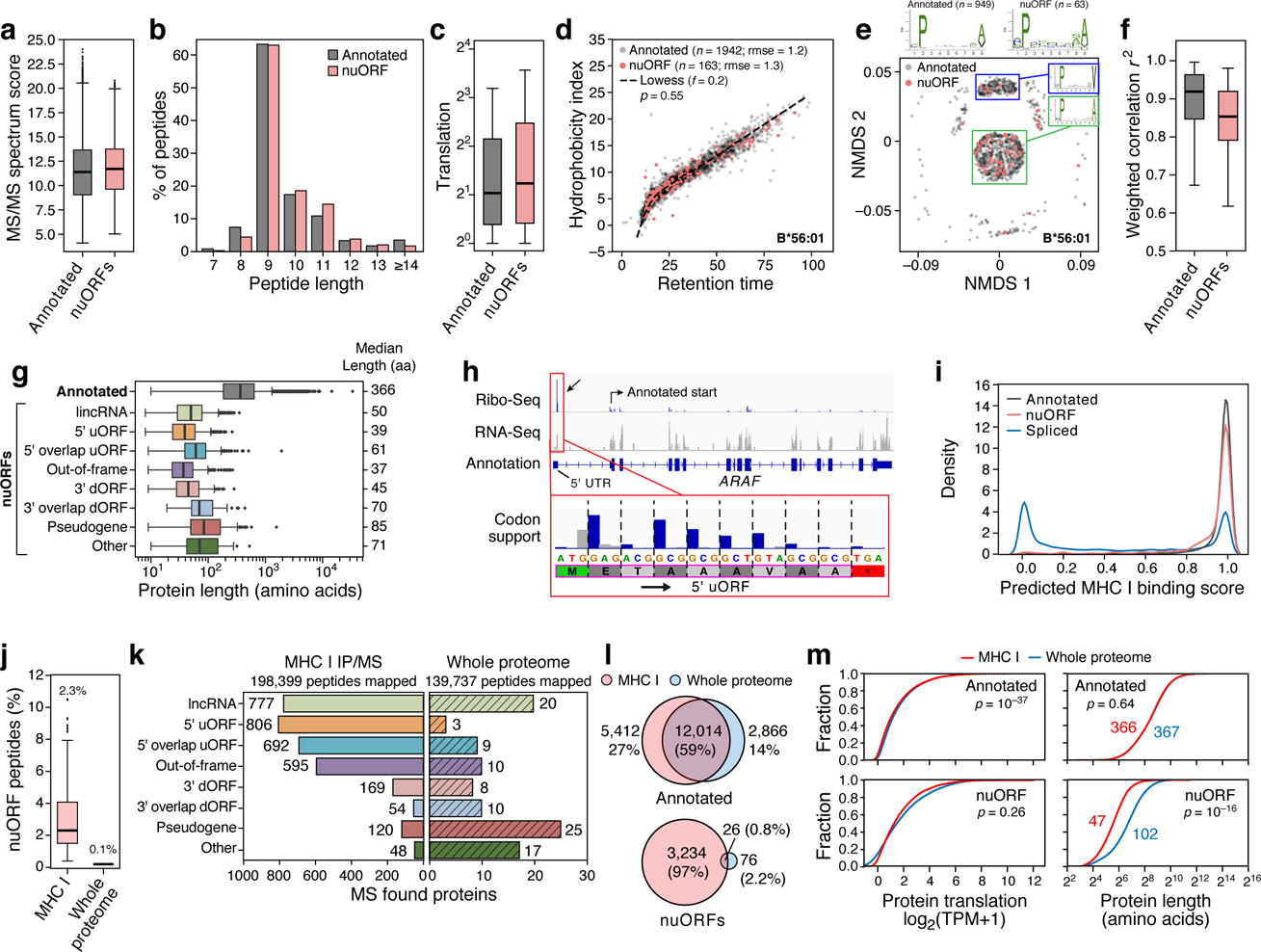
nuORFs peptides in the MHC I immunopeptidome have comparable biochemical properties to annotated ORFs. **a-g.** Comparable features of nuORFs and annotated peptides. **a.** LC-MS/MS Spectrum Mill identification score (y axis) for nuORF (pink) and annotated (grey) peptides (mean scores: 11.7 nuORF, 11.4 annotated; 2.4% to 3.8% increase, 95% CI). **b.** Distribution of detected peptide length (x axis) for nuORF (pink) and annotated (grey) peptides (median 9 AA for both). **c.** Ribo-seq translation levels (y axis, log2(TPM+1)) of annotated proteins (grey) and nuORFs (pink) in B721.221 cells (means: 1.6 annotated, 1.7 nuORF, 5.8% to 11.7% increase, 95% CI). **d.** Predicted hydrophobicity index (y axis) and retention time (x axis) of annotated (grey) and nuORF (pink) peptides for the HLA-B*56:01 sample. Dashed line: Lowess fit to the annotated peptides. **e.** Similar sequence motifs in nuORFs and annotated peptides. NMDS plot of all 9 AA peptides (dots) identified in HLA-B56:01 from nuORF (pink) or annotated ORFs (grey). Sequence motif plots shown for all annotated, all nuORF, and two marked clusters. **f.** Entropy weighted correlation (y axis) across all B721.221 HLA alleles between identified 9 AA annotated peptides and either down-sampled sets of annotated peptides, or nuORF peptides. **g.** nuORFs contributing peptides to the MHC I immunopeptidome are shorter than corresponding annotated proteins. Distribution of length (x axis) of different nuORF classes and annotated proteins (y axis) contributing peptides to the MHC I immunopeptidome. **h.** A 5’ uORF from *ARAF* detected in the MHC I immunopeptidome. Red box: magnified view of the 5’ uORF read coverage. Blue bars: in-frame reads, grey bars: out-of-frame reads. Magenta outline: LC-MS/MS detected peptide with periodicity plot showing strong read support for translation. **i.** Distribution of predicted MHC I binding scores for annotated peptides (grey), nuORF peptides (pink) and proteasomal spliced peptides from Faridi et al for 9 of our alleles (blue). j-m. nuORFs in the immunopeptidome have distinct characteristics compared to those in the whole proteome. **j.** Percent nuORFs (y axis) in immunopeptidome across 92 HLA alleles (pink) or of the whole proteome (grey). **k.** Number of nuORFs (x axis) of different categories (y axis) detected in the immunopeptidome (left) or the whole proteome (right). **l.** Proportion of all annotated ORFs (top) or nuORFs (bottom) detected in the whole proteome (blue), immunopeptidome (pink) or both (intersection) in B721.221 cells. **m.** Cumulative distribution function plots of Ribo-seq translation levels (left, x axis, log2(TPM+1)) or protein length (right, x axis) for annotated ORFs (top) or nuORFs (bottom) in MHC I immunopeptidome (red) or the whole proteome (blue). P-values: KS test. For all boxplots (A,C,F,H,J): median, with 25% and 75% (box range), and 1.5 IQR (whiskers) are shown.

### Short, overlapping nuORFs identified in the MHC I immunopeptidome

While 97% of MS-detected annotated ORFs could be predicted at the root, 33.8% (680) of the MS-detected nuORFs were exclusively predicted at the nodes in the leaves or clades (**Supplementary Figure 4a**), highlighting the heightened sensitivity of our hierarchical approach for identifying both sample-specific and shared nuORFs (**Figure 1c**, **Supplementary Figure 1a**).

For example, peptides derived from two overlapping 5’uORFs within the 5’UTR of the *LUZP1* transcript were detected by MHC I IP MS/MS in B721.221 cells across four different alleles (**Supplementary Figure 4b**). Due to the overlap of these ORFs, one was not predicted at the root, but was predicted in the CLL node, whereas the other 5’uORF was either translated at much lower levels or not at all.

Since many of the nuORFs that yielded MHC-I-presented peptides (2,093 of 3,261, 64%) overlapped with 5’UTRs and annotated ORFs, their identification from RNA-Seq alone would have been challenging, given their short length and proximity to, or overlap with, a longer annotated ORF, but they were readily identified by Ribo-seq (**Supplementary Figure 4b-d**). In fact, peptides from as many as three separate ORFs within one transcript were detected in the MHC I immunopeptidome. For example, for the *SOCS1* gene, an important modulator of interferon gamma and JAK-STAT signaling (Yoshimura, Naka, and Kubo 2007), peptides were identified matching the annotated protein, an internal out-of-frame nuORF (iORF) and a 5’ overlapping uORF (ouORF; **Supplementary Figure 4d**). Thus, nuORFs may be more readily expressed than previously anticipated, and not only generate peptides for MHC I presentation, but may also play other important roles in the cell.

As we previously reported for Ribo-seq predicted nuORFs (Ji et al. 2015), MHC I MS/MS-detected nuORFs were shorter than annotated ORFs (p < 10^-34^ across all nuORF types, *t*-test) (**Figure 2g**). Strikingly, the translated protein products of 26 nuORFs were exactly the same length as their corresponding MHC I-bound antigens, such that they should not require protease processing, as they are ready-made for MHC I presentation. For example, a peptide corresponding to an entire translated 5’ uORF from the 5’ UTR of *ARAF* is translated at a higher rate than the annotated *ARAF* protein in B721.221 cells (**Figure 2h**). The peptide matches the expected motif of HLA-B*45:01, where it was detected (**Supplementary Figure 5a**), and the LC-MS/MS spectra of the peptide closely support the sequence (**Supplementary Figure 5b**). Such short, abundant nuORF proteins may be presented on MHC I more quickly following translation than longer annotated proteins, which require protease processing.

### NuORF peptides explain some MS/MS spectra previously assigned to proteasomal spliced peptides

Proteasomal splicing of peptides has been proposed as a source of non-genomically encoded HLA class I antigens (Faridi et al. 2018; Liepe et al. 2016), but remains controversial as alternative interpretations for some of the underlying MS/MS spectra have been reported (Rolfs et al. 2019; Mylonas et al. 2018). For 9 of our previously published HLA class I-expressing monoallelic datasets (Abelin et al. 2017), our current data analysis found 308 nuORF-derived peptides, supported by Ribo-seq, that map to the same MS/MS spectra as 343 proposed spliced peptides (Faridi et al. 2018), in either of two scenarios: (1) for 98 cases, the nuORF-derived peptide sequence is identical to a proposed spliced peptide (**Supplementary Figure 6a,b**); or (2) for 210 cases, the partial sequence present in the MS/MS spectrum matched to a nuORF-derived peptide is also consistent with one or more different, yet similar, spliced peptide sequences (**Supplementary Figure 6c,d**). Notably, while 84% of nuORF peptides and 94% of annotated peptides had predicted MHC I binding scores over 0.8 (**Methods**), only 33% of proposed spliced peptides did (**Figure 2i**), consistent with reports that many spliced peptides were incorrectly identified (Rolfs et al. 2019; Mylonas et al. 2018). This suggests that stringent FDR thresholds and careful attention to sequence ambiguities resulting from *de novo* MS/MS interpretations are particularly warranted when interpreting the MS/MS spectra of peptides derived from noncanonical sources.

### NuORFs differ between the whole proteome and MHC I immunopeptidome

NuORFs were under-represented in whole proteome MS/MS analyses compared to the MHC I immunopeptidome, consistent with previous reports (Erhard et al. 2018; Raj et al. 2016). In the whole proteome of B721.221 cells, we identified 205 peptides from 102 nuORFs, representing only 0.1% of all peptides identified and >20-fold *fewer* peptides than in the MHC I immunopeptidome (**Figure 2j**). For example, we detected only 10 out-of-frame nuORFs and three 5’ uORFs in the whole proteome, compared to 595 and 806 such nuORFs in the MHC I immunopeptidome, respectively (**Figure 2k**). Additionally, while 59% of all detected annotated proteins were observed in both the MHC I immunopeptidome and in the whole proteome, only 0.8% of nuORFs were shared (**Figure 2l**). Despite comparable levels of translation between nuORFs detected on the MHC I and in the whole proteome (MHC I: 1.23, Whole: 1.42, p=0.26, KS test), the median length of nuORFs detected on the MHC I was far shorter than those detected in the whole proteome (**Figure 2m**, 47 vs. 102 amino acids, p < 10^-16^, KS test), suggesting a preference for presentation of shorter nuORFs on MHC I.

### NuORF identification in cancer MHC I immunopeptidomes

To further investigate nuORFs as a potential source of novel cancer antigens, we used nuORFdb to analyze the MHC I immunopeptidome of 10 cancer samples. On average, ∼1.5-2.2% of the immunopeptidome was assigned to nuORFs, across the melanoma, glioblastoma and CLL samples in nuORFdb (2.2%, n=3), as well as additional melanoma and glioblastoma samples (2.0%, n=5), and renal cell carcinoma and ovarian cancer samples not used to create the database (1.5%, n=2) (**Figure 3a**, **Supplementary Figure 7a**). Interestingly, compared to B721.221 cells, lncRNA nuORFs were less frequently observed across these primary human cancer samples (p = 10^-6^, ranksum test, see **Methods**), while 5’ uORFs were enriched (p = 0.05, rank-sum test, **Methods**) (**Supplementary Figure 7b-d**). NuORFs detected across various cancer samples were predicted from multiple nodes used in the generation of nuORFdb, with no single node able to account for all detected nuORFs in a given sample, highlighting the benefits of our hierarchical ORF prediction approach (**Figure 3b**). Importantly, nuORFdb helped detect MHC I presented peptides from translated nuORFs even in samples without any Ribo-seq data, albeit at lower proportion (**Figure 3a**). Overall, we detected peptides from 576 unique nuORFs of various types across all cancer immunopeptidomes (**Figure 3c**). More than half (50.6%) of the nuORFs were detected in more than one sample, demonstrating that they are not likely derived from random translation, but are translated recurrently across multiple samples (**Figure 3d**). As with B721.221 cells, nuORFs were under-represented in the whole proteome of a glioblastoma sample compared to the MHC I immunopeptidome (**Supplementary Figure 7e,f**), and those nuORFs detected in the whole proteome were significantly longer than those detected in the MHC I immunopeptidome (**Supplementary Figure 7g**, p = 10^-5^, KS test), with only 1% overlap between the two sets (**Supplementary Figure 7h**).

**Figure 3.**
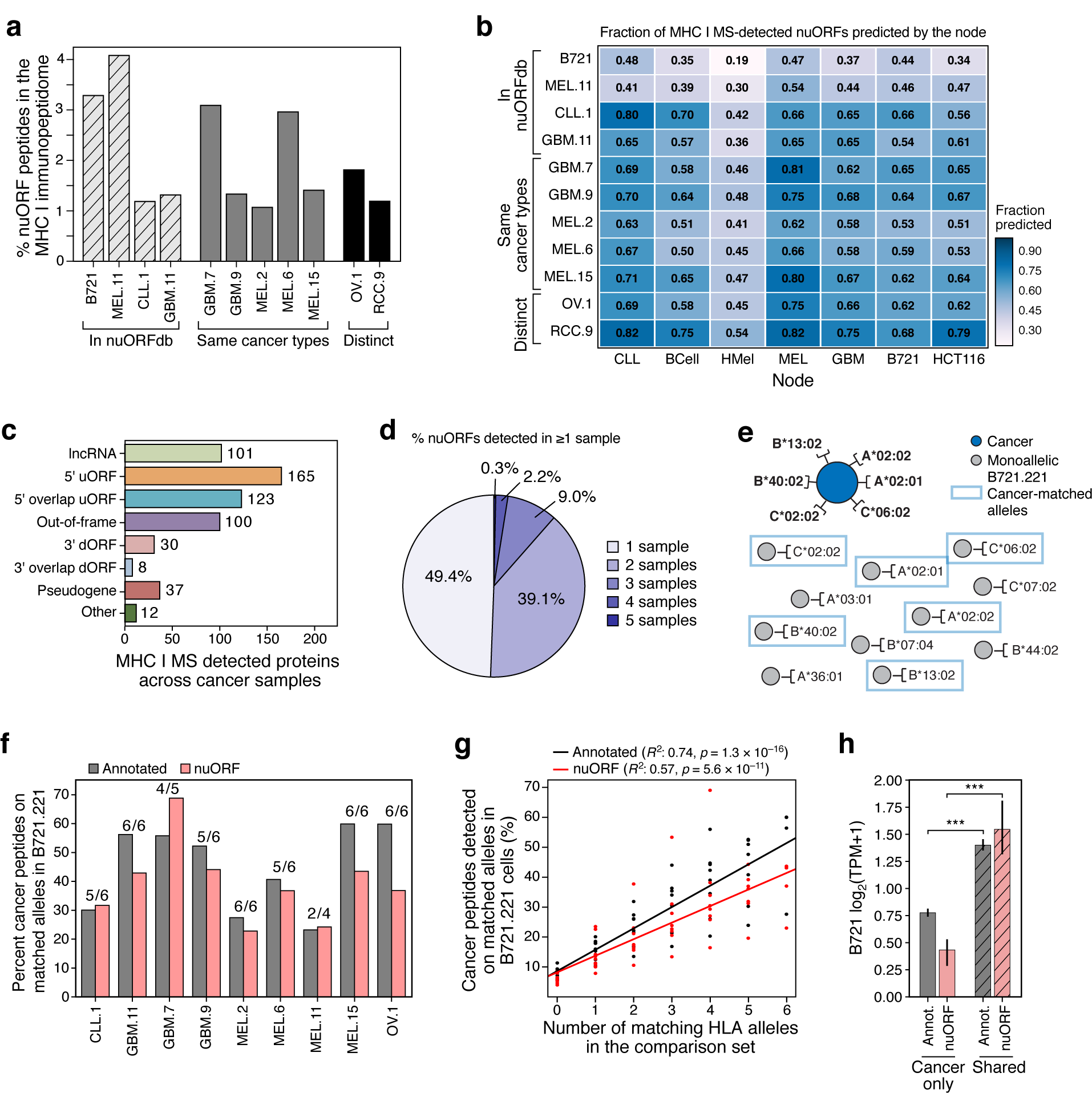
nuORF peptides in the MHC I immunopeptidome of cancer cells. **a-c.** nuORFdb allows detection of nuORFs in MHCI I immunopeptidome of samples and tumors types without prior Ribo-Seq data. **a.** Percent nuORF peptides detected in the MHC I immunopeptidome (y axis) from primary CLL, GBM, melanoma (MEL), ovarian carcinoma (OV), and renal cell carcinoma (RCC) (x axis). Hashed bars: Samples that contributed to nuORFdb. Grey bars: Same cancer types as in nuORFdb but from other patients. Black bars: Samples from tumor types not represented in nuORFdb. **b.** Fraction of MS/MS-detected nuORFs (colorbar) in each sample (rows) predicted by each node (columns). **c.** Number of nuORFs (x axis) of different types (y axis) identified in the MHC I immunopeptidome across 10 cancer samples. **d.** More than half of nuORFs are detected in more than one sample. Percent of nuORFs detected in one or more samples, including all cancer samples and B721.221 cells. e-h. Identical peptide sequences are presented on the same HLA alleles in cancer and in B721.221 cells. **e.** Approach to analyze peptide overlap between cancer samples and B721.221 cells expressing the same HLA alleles. Dark blue circle: cancer sample with 6 known HLA alleles. Grey circles: HLA mono-allelic B721.221 cells. Blue boxes: B721.221 cells used in the overlap analysis expressing cancer-matched HLA alleles. **f.** Percent of annotated (grey) and nuORF (pink) peptides (y axis) detected in cancer immunopeptidomes (x axis) that are also detected in HLA type-matched B721.221 samples. Number of available B721.221 sampled alleles over cancer sample’s known HLA alleles are shown above the bar. **g.** Percent of annotated (black) or nuORF (red) peptides (y axis) detected in cancer MHC I immunopeptidomes that are also detected in 6 B721.221 mono-allelic samples with variable numbers of HLA-matched samples (x axis). **h.** Median Ribo-seq translation levels (y axis, log2(TPM+1)) of annotated ORFs (grey) and nuORFs (pink) exclusive to cancer samples or also detected in B721.221 cells (hashed) (t-test, Annotated: p = 10-109, nuORF: p = 10-13). Error bars: 95% CI.

Identical peptide sequences were frequently detected in the cancer cells and in our HLA-matched B721.221 models (**Figure 3e,f**) for both annotated ORFs and nuORFs. The extent of overlap increased with the increase in the number of HLA alleles matching between B721.221 and the cancer cells (**Figure 3g**), for both annotated peptides (Pearson r^2^ = 0.74, p = 10^-16^) and nuORFs (r^2^ = 0.57, p = 10^-11^). Those ORFs that were detected in cancer cells but not in B721.221 cells had a lower level of translation in B721.221 cells, for both annotated ORFs (p = 10^-109^, t-test) and nuORFs (p < 10^-13^, t-test) (**Figure 3h**).

### NuORFs are sources of cancer antigens

Next, we estimated the extent to which nuORFs can serve as cancer antigens through either cancer-specific somatic mutations in nuORFs; or through cancer-specific translation (**Figure 4a**, **Supplementary Figure 8a**).

**Figure 4.**
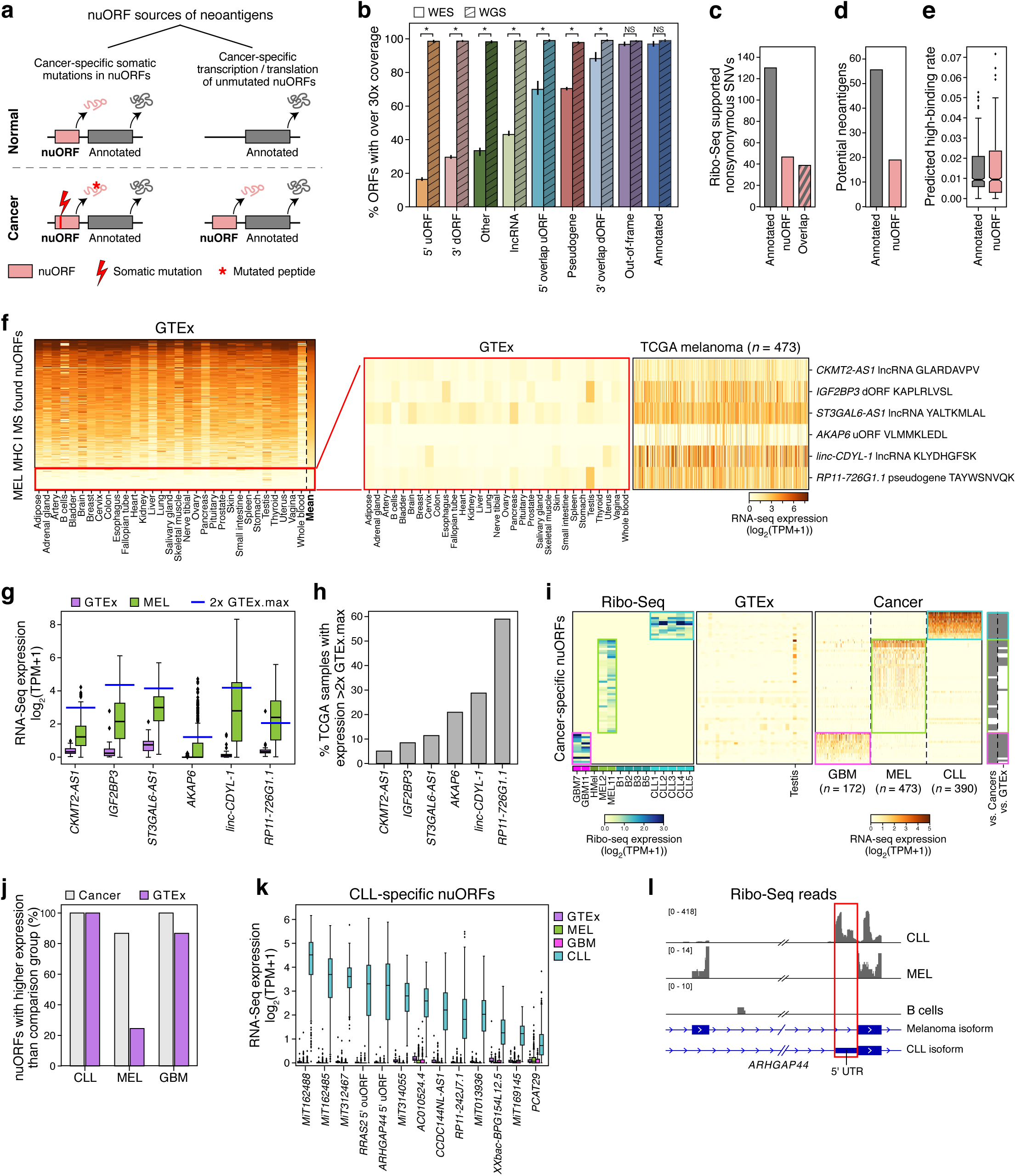
nuORFs expand the mutated and non-mutated antigen repertoire in cancer. **a.** Approaches to identify potential nuORF-derived neoantigens. **b-e.** Potential neoantigens from nuORFs with somatic mutations. **b.** Percent of ORFs with median ≥30× read coverage (y axis) by WES (n=18 samples: primary melanoma and GBM and matched normal) and WGS (n=2 samples: MEL11 and matched normal, hashed) for different types of ORFs (x axis) (*p < 0.01, t-test). Error bars: 95% CI. **c.** Number of Ribo-seq supported, non-synonymous SNVs (y axis) in MEL11 in annotated ORFs, nuORFs, or in both ORF types when they overlap. **d.** Number of high affinity (<500nM, netMHCpan v4.0) potential neoantigens (y axis) from annotated ORFs (grey) and nuORFs (pink) in MEL11. **e.** The rate of SNV-derived potential neoantigen peptides with high binding affinity (<500nM, netMHCpan v4.0) (y axis) from annotated ORFs (grey) and nuORFs (pink) across 1,170 netMHCpan v4.0 trained HLA alleles (means: 1.4% annotated, 1.6% nuORFs (0.1-0.3% higher, CI 95%)). For the boxplot, the median is shown, the 25% and 75% define the box range, and the whiskers go up to 1.5 IQR. f-h. MHC I MS/MS-detected nuORFs enriched in cancers may be potential sources of neoantigens. **f.** Expression level (log2(TPM+1)) of nuORFs (rows) detected in MHC I immunopeptidomes of 4 melanoma samples, ordered by mean expression (rightmost column) across all GTEx tissues (columns), except testis. Red box: nuORF at bottom 15% by mean expression (left), filtered for those expressed at least 2-fold higher than the maximum expression in GTEx in at least 5% of 473 melanoma samples in (TCGA) (right). **g.** Expression level (y axis, log2(TPM+1)) of melanoma-enriched, MS/MS-detected nuORFs in GTEx (purple) and TCGA melanoma (green) samples (x axis). Blue line: 2x highest GTEx expression (testis excluded). **h.** Percent of TCGA melanoma samples (y axis) with nuORF transcript (x axis) expression greater than 2x highest GTEx expression. **i-l.** nuORFs specifically translated in cancers as potential sources of neoantigens. **i.** Left: Ribo-seq translation levels (log2(TPM+1)) of nuORFs (rows) exclusively translated in GBM (pink box), melanoma (green box) or CLL (teal box) samples (columns, left), with median expression < 1 TPM across GTEx tissues (columns, middle) (testis excluded), and their expression (log2(TPM+1)) in respective cancer samples (columns, right). Far right: Significantly higher expression (grey, p < 0.0001, rank-sum test) in expected cancer type vs. the other cancer types or vs. GTEx expression. **j.** Percent of nuORFs (y axis) for each cancer type (x axis) with significantly higher expression (p < 0.0001, rank-sum test) in the expected cancer type than the other two cancer types (grey) or GTEx (purple) samples. **k.** Expression (y axis, log2(TPM+1)) of CLL-specific nuORFs (x axis) in CLL (teal), GBM (pink), melanoma (green), and GTEx (purple). **l.** CLL-specific *ARHGAP44* 5’ uORF (red box). Alternative transcript isoforms are translated in melanoma vs. CLL, and not translated in B cells. For all boxplots (E,G,K): median, with 25% and 75% (box range), and 1.5 IQR (whiskers) are shown.

For cancer-specific somatic mutations in nuORFs, we considered that WES is currently the standard approach to identify cancer-specific somatic mutations in annotated ORFs, yet it does not provide sufficient coverage to capture somatic mutations across various categories of nuORFs. While >99% of annotated ORFs had over the recommended 30X median coverage in WES, the coverage across nuORF types varied, and only 19.5% of 5’uORFs and 43% of nuORF-bearing lncRNAs had similar coverage in WES (**Figure 4b**, **Supplementary Figure 8b**, **Methods**). We therefore performed WGS, which provided at least 30X median coverage for over 98% of both annotated ORFs and nuORFs, across all types (**Figure 4b**, **Supplementary Figure 8b**).

To estimate the potential contribution of nuORFs with somatic mutations to the neoantigen repertoire in our systems, we focused our WGS analysis on a primary melanoma cell line (and matched PBMCs) previously characterized by WES, obtained from a patient who had received a personal neoantigen-targeting cancer vaccine (Ott et al. 2017); these cells were further profiled by Ribo-seq. The somatic mutation analysis by WGS closely recapitulated the original patient tumor genetics profiled by WES (**Supplementary Figure 8c**). We developed a computational pipeline to retrieve the Ribo-seq translation support for the mutant and wild-type alleles containing single nucleotide variants (SNVs) (**Supplementary Figure 8d**, **Methods**). We then selected the most likely neoantigens that could be derived from somatic variants in nuORFs and annotated ORFs, by prioritizing those that are: (1) mutated (by somatic mutation calling), (2) translated (by Riboseq coverage, in the mutant allele), and (3) predicted to be presented on MHC I (<500nM predicted MHC I binding affinity by netMHCpan 4.0 (Hoof et al. 2009)).

For this patient-derived melanoma sample, Ribo-seq supported the translation of a total of 217 SNVs, 22% of them exclusively in nuORFs (**Figure 4c**), with 19 of 75 (25%) of the mutated epitopes predicted to bind to autologous HLAs, derived from translated nuORFs (**Figure 4d**). Given the diversity of HLA alleles across the human population, based on the variants in this patient, we estimate that the rate at which mutations can generate antigens with high predicted MHC I binding affinity across alleles is 1.4% and 1.6% for annotated ORFs and nuORFs, respectively (**Figure 4e**, CI 95%: 0.1-0.3%). Altogether, these results suggest that nuORFs provide a sizeable additional source of potential neoantigens in cancer.

### Cancer-specific nuORF translation

Finally, we assessed the potential for neoantigen generation by cancer specific translation. To identify nuORFs translated in a melanoma-specific manner, we analyzed the 335 nuORFs detected in the MHC I immunopeptidomes from 4 melanoma samples (**Figure 4f**) and identified 6 high confidence melanoma-specific nuORF candidates. These were defined as those transcripts among the 335 nuORFs that were both lowly expressed across all healthy tissues (except the testis) in the Genotype-Tissue Expression (GTEx) collection of RNA-seq of healthy tissues (Consortium, G. TEx 2015) (**Figure 4f**, red box), and expressed at least 2-fold higher than the maximum GTEx expression in at least 5% of 473 melanoma samples in The Cancer Genome Atlas (TCGA) (Hutter and Zenklusen 2018; Blum, Wang, and Zenklusen 2018) (**Figure 4a,f**, **Methods**). Two of the six nuORFs, found in the *RP11-726G1.1* pseudogene and the *linc-CDYL-1* lncRNA, were highly overexpressed in 28% and 59% of TCGA melanoma samples respectively, suggesting potential shared candidate antigen targets across melanoma patients (**Figure 4g,h**).

We also used our Ribo-seq data to identify additional nuORFs that are not only translated in a cancer-specific manner (**Figure 4a,i**), but also lowly expressed by RNA-seq (TPM < 1, excluding testis) across healthy tissues in GTEx (**Figure 4i**). Indeed, many of the selected nuORFs had higher expression in the matched cancer type compared to healthy tissues and other cancer samples (p < 0.0001, **Figure 4i,j**). In particular, 13 nuORFs were strongly upregulated in CLL compared to GTEx and other cancer samples (**Figure 4k**). For example, we found a CLL-specific 5’uORF in the *ARHGAP44* locus, a gene which has been shown to be upregulated in CLL patients up to 10 years prior to diagnosis (Georgiadis et al. 2017). The *ARHGAP44* 5’ uORF is translated from a 5’UTR of a CLL-specific *ARHGAP44* transcript isoform, not expressed in healthy B cells and different from the isoform expressed in other tissues, such as melanoma (**Figure 4l**). Another CLL-specific 5’ouORF was detected in the *RRAS2* gene, which is upregulated in CLL patients with deletion in chromosome 13q (Rodríguez et al. 2012). Given the low frequency of somatic mutations in CLL (Rajasagi et al. 2014), these CLL-specific nuORFs could provide new antigenic targets for therapy.

We similarly identified several GBM and melanoma-specific nuORFs (**Supplementary Figure 9a,c**). In GBM, several nuORFs were translated from the *SOX2-OT* “noncoding” transcript (**Supplementary Figure 9a**) and a peptide (MIFESKTLF) derived from one of the *SOX2-OT* nuORFs was detected in the MHC I immunopeptidome of one of the GBM samples (**Supplementary Figure 9b**). *SOX2-OT*, annotated as a lncRNA, is frequently upregulated in GBM patients, and is essential for GBM tumorigenesis (Su et al. 2017). Given that *SOX2-OT* harbors several nuORFs specifically translated in GBM, further exploration of its role in GBM pathogenesis and potential immunogenicity is warranted.

## Discussion

Here, we combined Ribo-seq and MHC I immunopeptidome mass spectrometry analysis to identify thousands of nuORFs that were translated in healthy and cancer cells and contributed antigens to MHC I presentation. We detected both somatic mutations in nuORFs of cancer samples and nuORFs with tumor-specific translation, each expanding the pool of potential immunogenic cancer antigens.

Our comprehensive nuORF identification was enabled by our large Ribo-seq dataset collected across 29 samples of different tissue types, and our hierarchical ORF identification approach which leveraged this abundant data to identify nuORFs translated across tissues as well as in tissue- and sample-specific manner, thus constructing nuORFdb v1.0. The hierarchical nuORF identification pipeline leveraged the complementary power of tools with high nuORF discovery rates (RibORF (Ji et al. 2015)), particular power for identifying short overlapping out-of-frame nuORFs (PRICE (Erhard et al. 2018)), and recovery of cancer-specific nuORFs through *de novo* transcript assembly (MiTranscriptome (Iyer et al. 2015)). In this way, we maximized the number of translated nuORFs while maintaining a reasonable database size, making it a practical resource for routine use in MS studies.

Our extensive MHC-I immunopeptidome proteomics analysis allowed us to validate that nuORFs are translated and presented, and provided ground truth for improving prediction quality. The >6,000 nuORF peptides we identified with stringent criteria as presented on MHC I, have similar biochemical and biophysical characteristics to peptides derived from annotated proteins, and dramatically expand the number of nuORFs detected by mass spectrometry (Erhard et al. 2018; Martinez et al. 2019; Ma et al. 2016, 2014), particularly in primary cancer samples. As 50.6% of nuORFs were detected in more than one MS sample, they are recurrently translated and MHC I-presented, with identical peptide sequences frequently detected in patient-derived cancer cell lines and in B721.221 mono-allelic cells expressing matching alleles, highlighting the robustness of nuORF prediction and the dependency of MHC I presentation on the HLA allele expressed in a given sample. Notably, the annotations in nuORFdb provide better context for interpretation of MHC I immunopeptidome data, favoring for example nuORF explanations of many peptide sequences previously postulated to be derived from proteasomal splicing (Faridi et al. 2018). While nuORFdb is likely not fully saturated, it can already be used to identify nuORFs in tissue types not yet profiled by Ribo-seq.

Far fewer nuORFs were detected in whole proteome vs. MHC I immunopeptidomes analysis, despite their similar translation levels, consistent with previous reports that nuORFs are rarely detected in whole proteomes (Erhard et al. 2018; Raj et al. 2016). NuORFs detected on MHC I were substantially shorter than those detected in the whole proteome, suggesting that shorter nuORFs may be more likely presented on MHC I. Interestingly, 26 of the nuORFs encoded peptides exactly the same length as their observed MHC I-bound antigens, requiring no additional post-translational processing prior to presentation. Translation of such peptides may result in a more rapid MHC I presentation and recognition by the immune system, thus providing a faster readout of cellular state. To our knowledge, these are the first examples of very short nuORFs that are translated and detected on MHC I without post-translational processing.

While the primary biological function of some nuORFs may be to become antigens that trigger an immune response, it is likely that others might have additional biological functions. In particular, we have detected peptides from 318 nuORFs in transcripts currently annotated as lincRNAs, which should be prioritized in future perturbation studies. In other examples, the hierarchical nuORF identification approach detected overlapping, out-of-frame nuORFs, encoded by the same gene, such as *SOCS1*, involved in the interferon gamma signaling pathway (Yoshimura, Naka, and Kubo 2007). In our MHC I immunopeptidome data, we identified peptides derived from 3 overlapping proteins with completely different amino acid sequences, all encoded by the *SOCS1* gene. They included the annotated SOCS1 protein, a short out-of-frame iORF, and another short 5’ ouORF, overlapping, but out-of-frame with the 5’ end of the annotated ORF. Given the important biological function of the annotated SOCS1 protein, the cellular roles of *SOCS1*-derived nuORFs beg investigation, as do the dynamics of their translation.

Both somatic mutations in nuORFs and cancer specific translation of nuORFs can expand the neoantigen repertoire. In melanoma, we identified 25% more potential neoantigens derived from translated nuORFs containing somatic mutations. While WGS successfully captured variants across all nuORF types in nuORFdb, WES frequently exhibited insufficient coverage, in particular, for nuORFs in 5’ and 3’ UTRs and lncRNAs. Expanding WES panels to include the UTRs of protein-coding transcripts harboring nuORFs could extend clinical access to an expanded pool of potential neoantigens. Among the nuORFs transcribed and translated in cancer-specific manner in melanoma, GBM or CLL, were a 5’ uORF in *ARHGAP44*, a 5’ ouORF in *RRAS2*, and nuORFs in *SOX2-OT* lincRNA, each derived from a gene involved in cancer biology (Georgiadis et al. 2017; Rodríguez et al. 2012; Su et al. 2017). Multiple additional cancer-specific nuORFs were derived from novel transcripts (Iyer et al. 2015). These cancer-specific nuORFs can be potential sources of cancer antigens or carry important biological functions.

In summary, nuORFs are prevalent, translated, and contribute ∼2% of the antigen repertoire presented on MHC I in cancer cells. NuORFs can harbor somatic mutations, and can be translated in a cancer-specific manner, providing an expanded source of potential immunogenic targets. Additionally, the large number of such varied types of nuORFs being translated raises the question of what biological functions they have beyond immune presentation. Our data suggest that nuORFs should be considered when exploring the immunopeptidome, and that they reveal attractive candidates for immunotherapeutic targeting and biological investigation.

## Author contributions

T.O. and A.R. conceived the study. D.B.K., S.A.C., C.J.W., N.H. and A.R. directed the overall study design. T.O., E.C. and Y.T.C. generated Ribo-seq libraries. T.O. and T.L. performed Riboseq analysis. S.K., K.R.C., T.O., T.L., S.S., C.R.H., H.K. and A.A. generated the MS data and performed the associated data analysis. B.A.K. provided CLL RNA-seq data. F.A. performed GTEx, TCGA and CLL RNA-seq alignment and quantification under G.G. guidance. B.L. performed whole genome sequencing analysis. D.B.K. and P.L. generated the single-HLA allele cell lines. D.B.K., G.O. and C.J.W. provided the patient-derived tumor cell lines. P.B. provided CLL samples. P.B., W.Z. and D.B.K. prepared PBMCs and B cells from CLL patients and healthy donors. I.J. performed conservation analysis under M.K. guidance. Z.J. and S.A.S. provided computational support. T.O., T.L., K.R.C., S.K., S.S., D.B.K., S.A.C., C.J.W., and A.R. wrote the manuscript, with contributions from all co-authors.

## Acknowledgements

We thank Kirk Gosik and Rebecca Herbst for their help with the statistical analysis. We thank Doris Fu for her help with the NMDS analysis. We thank Eran Hodis and John Kwon for providing cultured primary melanocytes. We thank Keith L. Ligon for providing the GBM cell line. We thank Leslie Gaffney for help with figure preparation. Work was supported by the Klarman Cell Observatory and HHMI (A.R.), NIH grants NCI-1R01CA155010-02 (to C.J.W.), NHLBI-5R01HL103532-03 (to C.J.W.), NIH/NCI R21 CA216772-01A1 (to D.B.K.), NCI-SPORE-2P50CA101942-11A1 (to D.B.K), NHGRI T32HG002295 and NIH/NCI T32CA207021 (to S.S.), NCI R50CA211482 (to S.A.S.), NHGRI U41HG007234 and R01 HG004037 (to I.J.), NCI Clinical Proteomic Tumor Analysis Consortium grants NIH/NCI U24-CA210986 and NIH/NCI U01 CA214125 (to S.A.C.) and NIH/NCI U24CA210979 (to D.R. Mani and Gad Getz). This work was supported in part by The G. Harold and Leila Y. Mathers Foundation and the Bridge Project, a partnership between the Koch Institute for Integrative Cancer Research at MIT and the Dana-Farber/Harvard Cancer Center. C.J.W. is a scholar of the Leukemia and Lymphoma Society, and is supported in part by the Parker Institute for Cancer Immunotherapy. S.K. is a Cancer Research Institute/Hearst Foundation fellow. T.O. is a Leukemia and Lymphoma Society Fellow. B.A.K. is supported by a long-term EMBO fellowship (ALTF 14-2018). P.B. is supported by an Amy Strelzer Manasevit Grant and an American Society of Hematology Scholar Award. G.O. is supported by a postdoctoral fellowship sponsored by the American-Italian Cancer Foundation.

## Methods Cell cultures

A375 cells were cultured in DMEM media (Gibco), supplemented with 5% fetal bovine serum (FBS). HCT116 cells were cultured in McCoy’s 5A Medium (Thermo Fisher Scientific), supplemented with 5% FBS.

### Generation of HLA mono-allelic B721.221 cells

The HLA mono-allelic cell lines were generated as previously described (Abelin et al. 2017; Sarkizova et al. 2019). Briefly, single HLA allele-expressing cDNA vectors in a pcDNA-3 backbone were ordered from GenScript^TM^. The HLA class I deficient B721.221 cell line was transfected with the HLA allele expression vectors using lipofectamine, as described previously (Abelin et al. 2017). Cell lines with stable surface HLA expression were generated first through selection using 800µg/ml G418 (Thermo Fisher Scientific), followed by enrichment of HLA positive cells through up to 2 serial rounds of fluorescence-activated cell sorting (FACS) and isolation using a pan-HLA antibody (W6/32; Santa Cruz) on a FACSAria II instrument (BD Biosciences).

### Primary human cells and generation of cancer cell lines

All human tissues were obtained following informed consent through DFCI or Partners Healthcare approved IRB protocols. Conditions for growth and *in vitro* propagation of melanoma and GBM tumor cell lines were described previously (Ott et al. 2017; Keskin et al. 2019). PBMCs from fresh healthy donor whole blood were isolated using Ficoll density gradient medium. CD19^+^ B cells were isolated using EasySep Human CD19 Positive Selection Kit, obtaining between 25 and 54 million B cells per donor. For fresh CLL samples, PBMCs were isolated using Ficoll density gradient medium, enriched for CD19 positive CLL tumor cells and were used in IP/MS analysis and Ribo-seq. For cryopreserved CLL samples, live cells were isolated with an OptiPrep density gradient medium. Surgically resected clear cell renal cell carcinoma (ccRCC) tissue was mechanically dissociated with scalpels, and then enzymatically dissociated using a mixture of collagenase D (Roche), Dispase (STEMCELL Technologies), and DNase I (New England BioLabs) at room temperature, and filtered through a 100 micron cell strainer using the sterile plunger of a syringe. Red blood cells were lysed using ammonium-chloride-potassium buffer (Gibco). The cell suspension was stained for viability (Zombie Aqua; BioLegend), anti-CD45 (BV605; BD Biosciences), and anti-carbonic anhydrase IX (PE; R&D Systems). Viable, CD45^-^, CAIX^+^ tumor cells were isolated by FACS (BD FACSAria II cell sorter; BD Biosciences). Cells were cultured in a specialized growth medium consisting of OptiMEM GlutaMax media (Gibco), 5% fetal bovine serum, 1mM sodium pyruvate (Gibco), 100 units/mL penicillin and streptomycin, 50 micrograms/mL gentamicin, 5 micrograms/mL insulin (Sigma), and 5 ng/mL epidermal growth factor (Sigma). Following successive passages, CAIX expression was confirmed by flow cytometry (anti-CAIX, PE-conjugated; R&D Systems) and by immunohistochemical analysis of a cell pellet.

Ovarian cancer patient-derived cells were propagated within a xenograft model, which was generated by serial passaging of tumor cells from a patient with advanced ovarian cancer. These cells originated from solid tumors that were injected orthotopically in the abdominal cavity in NOD-SCID mice (8-week old, Jackson labs). Tumor growth was monitored weekly by observing mice for signs of abdominal distension. Cells were harvested 4 months after initial injection and cryopreserved.

### Ribosome profiling

Ribosome profiling was performed according to the manufacturer’s protocol (TruSeq Ribo Profile - RPHMR12126, Illumina, discontinued), with the following modifications. For adherent cell lines (melanoma, primary melanocytes, HCT116, A375), culture media was removed, cells were washed with ice-cold PBS containing cycloheximide (0.1mg/ml) and lysed in the Lysis Buffer according to the Illumina protocol. For suspension cell lines and primary blood samples, cells were spun 1,000rpm for 5 minutes, washed once with ice-cold PBS containing cycloheximide (0.1mg/ml) and lysed in the Lysis Buffer. To perform Ribo-seq on small samples, such as primary B cells and melanocytes, cells were lysed in 200 µl of lysis buffer, such that the entire lysate could be used in library preparation. Ribosomes containing ribosome-protected mRNA fragments (RPFs) were enriched using MicroSpin S-400 columns (GE Healthcare, catalog # 27-5140-01). Ribo-zero rRNA Removal Kit (Illumina, MRZH11124, discontinued) was used to deplete rRNA from RPFs. The RPF sample was loaded on a 15% urea-polyacrylamide gel. Samples were eluted from the gel overnight at 4°C. Subsequently, end repair, adapter ligation and reverse transcription were carried out according to the manufacturer’s protocol. For the cDNA gel purification, the reverse transcription reaction was loaded on a 10% urea-polyacrylamide gel. The samples were eluted from the gel overnight at room temperature. Subsequently, RPFs were circularized and 5 µl of circDNA was used for library amplification. The number of amplification cycles was determined based on the observed sample quality and expected yield, but usually ranged between 8 and 10 cycles. Following amplification, the library was gel-purified using 4% E-Gel EX Agarose Gel (ThermoFisher G401004) and Zymoclean Gel DNA Recovery Kit (Zymo Research D4007), with 4 volumes of ADB buffer to accommodate 4% agarose gel. The resulting libraries were analyzed for quality using Agilent Bioanalyzer 2100 and sequenced for 51 cycles on the Illumina NextSeq platform, using NextSeq 500 high output kit, V2, 75 cycles.

### Ribo-seq data pre-processing

To process RPF sequencing reads, Illumina adapters were removed using fastx_clipper from the FASTX-Toolkit. Ribosomal RNA and tRNA were removed using Bowtie version 1.0.0 (Langmead et al. 2009). Remaining reads were aligned to the genome (hg19 / GRCh37) and transcriptome using STAR version 2.5.3a (Dobin et al. 2013) (--alignIntronMin 20 -- alignIntronMax 100000 --outFilterMismatchNmax 1 --outFilterType BySJout -- outFilterMismatchNoverLmax 0.04 --twopassMode Basic). For the transcriptome annotation, a combination of GENCODE v26lift37 transcriptome annotation was combined with transcripts annotated as *tstatus “unannotated”* from MiTranscriptome annotation (Iyer et al. 2015). To determine the RPF library quality, trinucleotide codon periodicity was plotted using RibORF readDist script (Ji et al. 2015) against annotated protein-coding ORFs (GENCODE v26lift37). Only samples and read lengths that showed clear trinucleotide periodicity were used for subsequent ORF predictions.

### Hierarchical prediction of translated open reading frames across tissues

In order to maximize the detection of translated ORF and overcome noise from overlapping ORFs expressed in different tissues, we performed hierarchical ORF predictions using RibORF (Ji et al. 2015) and PRICE (Erhard et al. 2018), as follows. **For RibORF**, only read lengths that showed clear trinucleotide periodicity were used for ORF predictions. RibORF offsetCorrect script was used to correct the RPF offsets for each read length.

As input, for the transcriptome reference, GENCODE v26lift37 transcriptome annotation was combined with transcripts annotated as *tstatus “unannotated”* from MiTranscriptome annotation (Iyer et al. 2015). From this custom transcriptome reference, all possible ORFs with NTG start codons and TAA/TGA/TAG stop codons were identified using Rp-Bp prepare-rpbp-genome script (Malone et al. 2017). For the GENCODE ORF search, Rp-Bp reported the following ORF types based on the annotation of the transcript and the location of the ORF within the transcript:

- *canonical*: identical to a protein-coding ORF annotated in the GENCODE reference.
- *canonical_extended*: Predicted start is 5’ extended relative to a protein-coding ORF annotated in the GENCODE reference.
- *canonical_truncated*: Predicted start codon is 3’ downstream of the annotated start codon in the GENCODE reference.
- *five_prime*: ORF entirely contained in the 5’ UTR of a protein-coding transcript.
- *five_prime_overlap*: ORF with a start codon in the 5’ UTR of a protein-coding transcript, and a stop codon within an annotated ORF, out-of-frame relative to the annotated ORF.
- *three_prime*: ORF entirely contained in the 3’ UTR of a protein-coding transcript.
- *three_prime_overlap*: ORF with a start codon within an annotated ORF, and the stop codon in the 3’ UTR, out-of-frame relative to the annotated ORF.
- *within*: entirely contained within, but out of-of-frame relative to an annotated ORF.
- *noncoding*
- *suspect*

Those ORFs annotated as *noncoding* or *suspect* by Rp-Bp were re-annotated based on the metadata column in the GENCODE GTF. The ORFs derived from transcripts containing ‘linc’ or ‘pseudo’ in the metadata column were annotated as *noncoding_lincRNA* or *noncoding_pseudogene* respectively. Otherwise, they were re-annotated as *noncoding_other*. For the MiTranscriptome transcripts, Rp-Bp reported all ORFs as either *noncoding* or *suspect*. Subsequently, the ORF types were re-annotated as *noncoding_mi_lincRNA* or *noncoding_mi_tucp* based on the transcript type annotated in the MiTranscriptome GTF as either *tcat “lncrna”* or *tcat “tucp”* respectively. After running RibORF, ORFs with a score > 0.7 were retained. If multiple ORFs on the same transcript shared a common stop codon, the longest ORF was selected.

### Hierarchical ORF prediction using RibORF

Offset-corrected SAM files across samples were combined at each clade and at the root (Supplementary Fig. 1a). For the ORFs predicted at the *root*, we retained predicted ORFs with at least 2 reads in-frame and a RibORF score > 0.7. For ORFs predicted at the cladees and leaves (**Supplementary Fig. 1a**) we retained predicted ORFs with at least 2 reads and score > 0.9, or at least 250 reads and score > 0.7.

**For PRICE**, we ran the PRICE pipeline (Erhard et al. 2018) on unprocessed fastq.gz files of the samples that had clear tri-nucleotide periodicity (as determined by RibORF above) with the same reference transcriptome as for RibORF. The pipeline handled adapter trimming, rRNA and tRNA removal, offset correction and ORF prediction. Unique .cit files were generated for each sample. For the **hierarchical ORF prediction using PRICE**, gedi MergeCIT was used to merge samples by tissue type at each clade and at the root. gedi Price -fdr 1 was used to predict translated ORFs.

The PRICE ORF annotation types (Erhard et al. 2018) and https://github.com/erhard-lab/gedi/wiki/Price include the following:

- CDS: ORF is exactly as in the annotation
- Ext: ORF contains a CDS, ending at its stop codon
- Trunc: ORF is contained in a CDS, ending at its stop codon
- Variant: ORF ends at a CDS stop codon, but is neither Ext nor Trunc
- uoORF: ORF starts in 5’-UTR, ends within a CDS
- uORF: ORF starts and ends in 5’-UTR
- iORF: ORF is contained within a CDS
- dORF: ORF ends in 3’-UTR
- ncRNA: ORF is located on non-coding transcript
- intronic: ORF is located in an intron
- orphan: Everything else

### Generating nuORFdb v1.0

FASTA files of ORFs predicted across tissues by RibORF and PRICE were combined, and those ORFs entirely contained within other predicted ORFs at the protein level were removed. Predicted ORFs over 21 nucleotides long were retained for the downstream analysis, and translated in the single frame determined from Ribo-seq periodicity. After merging the predictions from RibORF and PRICE, the nuORFdb contains the ORF types from both prediction tools, as described above. To improve annotations, for nuORFs in categories ncRNA, noncoding_other, orphan, and Variant, we identified their transcript_type annotated in the GENCODE GTF metadata and generated the nuORF Refined type. In order to unify the different terms for the same concept we subsequently merged the refined ORF types according to the specifications of biotypes in Ensembl (https://useast.ensembl.org/info/genome/genebuild/biotypes.html), generating an ORF type mapping table, where MergedType is used in Supplementary Fig. 3a, and PlotType is used in the rest of the figures, also shown in Fig.1e.

### HLA-peptide immunoprecipitation, sequencing by tandem mass spectrometry, and peptide identification

Soluble lysates from up to 50 million HLA expressing B721.221 cells or 0.1 to 0.2g cancer cells were immunoprecipitated with W6/32 antibody (sc-32235, Santa Cruz) as described previously (Abelin et al. 2017; Sarkizova et al. 2019). 10 mM iodoacetamide was added to the lysis buffer to alkylate cysteines for 71 alleles and 10 tumor samples. Peptides of up to three IPs were combined, acid eluted either on StageTips or SepPak cartridges (Bassani-Sternberg et al. 2016), and analyzed in technical duplicates using LC-MS/MS. Peptides were resuspended in 3% ACN, 5% FA and loaded onto an analytical column (20-30 cm, 1.9 µm C18 Reprosil beads (Dr. Maisch HPLC GmbH), packed in-house PicoFrit 75 µm inner diameter, 10 µm emitter (New Objective)). Peptides were eluted with a linear gradient (EasyNanoLC 1000 or 1200, ThermoFisher Scientific) ranging from 6-30% Buffer B (either 0.1% FA or 0.5% AcOH and 80% or 90% ACN) over 84 min, 30-90% B over 9 min and held at 90% Buffer B for 5 min at 200 nl/min. During data dependent acquisition, peptides were analyzed on a QExactive Plus (QE+), QExactive HF (QE-HF) or Fusion Lumos (ThermoFisher Scientific). Full scan MS was acquired at a resolution of 70,000 (QE+) or 60,000 (QE-HF and Lumos) from 300-1,800 m/z or 300-1,700 m/z (Lumos). AGC target was set to 1e6 and 5 msec max injection time for QE type instruments and 4e5 and 50 ms for Lumos. The top 10 (Lumos, QE+), 12 (QE+), 15 (QE-HF) precursors per cycle were subjected to HCD fragmentation at resolution 17,500 (QE+) or 15,000 (QE-HF, Lumos). The isolation width was set to 1.7 m/z with a 0.3 m/z offset for QE and 1.0 m.z and no offset for Lumos, the collision energy was set to optimal for the instrument used ranging from 25 to 30 NCE, AGC target was 5E4 and max fill time 120 ms (QE+ and Lumos) or 100 ms (QE-HF). For Lumos measurements, precursors of 800-1700 m/z were also subjected to fragmentation if they were singly charged. Dynamic exclusion was enabled with a duration of 15 sec (QE+), 10 secs (QE-HF) or 5 sec (Lumos).

### HLA peptide identification using Spectrum Mill

Mass spectra were interpreted using the Spectrum Mill software package v6.1 pre-Release (Agilent Technologies, Santa Clara, CA). MS/MS spectra were excluded from searching if they did not have a precursor MH+ in the range of 600-4000, had a precursor charge >5, or had a minimum of <5 detected peaks. Merging of similar spectra with the same precursor m/z acquired in the same chromatographic peak was disabled. Prior to searches, all MS/MS spectra had to pass the spectral quality filter with a sequence tag length >2 (*i.e.*, minimum of 4 masses separated by the in-chain masses of 3 amino acids).

MS/MS spectra were searched against the 323,848 protein sequences in nuORFdb v1.0 appended to a base reference proteome containing all UCSC Genome Browser genes with hg19 annotation of the genome and its non-redundant protein coding transcripts (52,788 entries) as well as 264 common laboratory contaminants, including proteins present in cell culture media and immunoprecipitation reagents. MS/MS data from patient derived cell lines was analyzed in the same way, except that the sequence database was revised with further inclusion of patient-specific somatic mutations.

MS/MS search parameters included: no-enzyme specificity; fixed modification: cysteinylation of cysteine; variable modifications: carbamidomethylation of cysteine, oxidation of methionine, and pyroglutamic acid at peptide N-terminal glutamine; precursor mass tolerance of ±10 ppm; product mass tolerance of ± 10 ppm, and a minimum matched peak intensity of 30%. Variable modification of carbamidomethylation of cysteine was only used for HLA alleles that included an alkylation step. Peptide spectrum matches (PSMs) for individual spectra were automatically designated as confidently assigned using the Spectrum Mill auto-validation module to apply target-decoy based FDR estimation at the PSM level of <1% FDR. Peptide auto-validation was done separately for each HLA allele with an auto thresholds strategy to optimize score and delta Rank1 – Rank2 score thresholds separately for each precursor charge state (1 through 4) across all LC-MS/MS runs for an HLA allele. Score threshold determination also required that peptides had a minimum sequence length of 7, and PSMs had a minimum backbone cleavage score (BCS) of 5. BCS is a peptide sequence coverage metric and the BCS threshold enforces a uniformly higher minimum sequence coverage for each PSM, at least 4 or 5 residues of unambiguous sequence. The BCS score is a sum after assigning a 1 or 0 between each pair of adjacent AA’s in the sequence (max score is peptide length-1). To receive a score, cleavage of the peptide backbone must be supported by the presence of a primary ion type for HCD: b, y, or internal ion C-terminus (*i.e.*, if the internal ion is for the sequence PWN then BCS is credited only for the backbone bond after the N). The BCS metric serves to decrease false-positives associated with spectra having fragmentation in a limited portion of the peptide that yields multiple ion types. PSMs were consolidated to the peptide level to generate lists of confidently observed peptides for each allele using the Spectrum Mill Protein/Peptide summary module’s Peptide-Distinct mode with filtering distinct peptides set to case sensitive. A distinct peptide was the single highest scoring PSM of a peptide detected for each allele. MS/MS spectra for a particular peptide may have been recorded multiple times (*e.g.*, as different precursor charge states, from replicate IPs, from replicate LC-MS/MS injections). Different modification states observed for a peptide were each reported when containing amino acids configured to allow variable modification; a lowercase letter indicates the variable modification (C-cysteinylated, c-carbamidomethylated).

In cases where a spectrum could be matched to multiple proteins due to shared peptide sequences, the Spectrum Mill output was revised so that the primary protein assignment for a spectrum was determined using the following decision tree, in order of diminishing assignment priority: Contaminants → annotated proteins → nuORFs. In cases where a spectrum could be matched to multiple annotated proteins, priority was given to the more highly translated one based on Riboseq TPM. In cases where a spectrum could be matched to multiple nuORFs, priority was given to the more highly translated based on Ribo-seq TPM. In case of equal Ribo-seq TPM, the primary assignment was randomly selected.

### FDR filtering of nuORF-derived peptides

Applying the same aggregate FDR threshold to the combination of peptides observed for both annotated ORFs and nuORFs resulted in a much higher FDR for nuORFs (4.6%) than for annotated ORFs (1%), which was as high as 14% for certain nuORF categories, such as 3’ overlapping dORFs (Supplementary Fig. 2c,d). We therefore introduced more stringent filtering for nuORF peptides (Supplementary Fig. 2e,f), to retain only the 6,501 which achieved <1% peptide-level FDR (Supplementary Fig. 2a-d,g).Spectra were removed based on fixed thresholds for 4 Spectrum Mill MS/MS scoring metrics: score, backbone cleavage score (BCS), BCS%, and percent scored peak intensity, defined as follows:

- Score: the primary score based on assignment of the full range of ion types (y, b, a, internal and neutral losses of NH_3_ and H_2_O) to peaks in a spectrum.
- Backbone cleavage score (BCS): absolute peptide sequence coverage metric described above
- BCS %: BCS normalized for peptide length, 100 * BCS / (sequence length - 1)
- Percent scored peak intensity: Percent of product ion intensity in an MS/MS (after peak detection) that is matched to a scored ion type.

NuORFs across all 92 alleles were binned by ORF type. FDRType column and integer thresholds were calculated per bin to maximize retained spectra with an FDR less than 1% (Supplementary Fig. 2c,d). Maximal thresholds were calculated using a grid search of integer threshold values encompassing the empirically observed values. Specifically, we identified the combination of lowest values across the 4 scoring metrics that resulted in FDR < 1% for each ORF type bin.

### Peptide spectrum matching with proposed splice peptides

For 9 of our previously published monoallelic datasets (A*02:03, A*02:04, A*02:07, A*03:01, A*24:02, A*31:01, A*68:02, B*44:02, B*51:01) (Abelin et al. 2017) that have been proposed to contain proteasomal spliced peptides (Faridi et al. 2018) we reanalyzed the data to examine if nuORF derived peptides could be better explanations for the spectra matched to proposed splice peptides. Since we were unable to determine which spectra were matched to individual spliced peptides for our datasets, we took the proposed spliced peptides in the supplemental tables from Faridi et al 2018 and appended them to our nuORFdb/Reference proteome database and repeated the analysis of the spectra for these 9 alleles using the process described above.

### Peptide hydrophobicity index calculation

Hydrophobicity index was predicted using SSRCalc (Krokhin et al. 2004), http://hs2.proteome.ca/SSRCalc/SSRCalcQ.html). Modification of cysteine was checked for alleles B5601 and A7401. For A0201 and C0304 free Cysteine was specified.

### Whole proteome analysis and interpretation

Protein expression of the B721.221 and GBM H4152-BT145 cell lines was assessed as described previously (Mertins et al. 2018). Briefly, cell pellets of B721.221 cells expressing A*03:01, B*55:01 and C*07:01, as well as pellets of GBM6 with and without IFNγ treatment were lysed in 8M Urea and digested to peptides using LysC and Trypsin (Promega). B721 analysis was performed label free with a 1:1:1 mix using 100 µg each of the three monoallelic cell lines. For GBM, 100 µg peptides were labeled with TMT6 reagents (Thermo Fisher) 126 (untreated) and 127 (IFNγ) and then pooled for subsequent fractionation and analysis. Pooled peptides were separated into 24 fractions using offline high pH reversed phase fractionation. 1 µg per fraction was loaded onto an analytical column (20-30 cm, 1.9 µm C18 Reprosil beads (Dr. Maisch HPLC GmbH), packed in-house PicoFrit 75 µM inner diameter, 10 µM emitter (New Objective)). Peptides were eluted with a linear gradient (EasyNanoLC 1000 or 1200, Thermo Scientific) ranging from 6-30% Buffer B (either 0.1%FA or 0.5% AcOH and 80% or 90% ACN) over 84 min, 30-90% B over 9 min and held at 90% Buffer B for 5 min at 200 nl/min. During data dependent acquisition, peptides were analyzed on a Fusion Lumos (Thermo Scientific). Full scan MS was acquired at a 60,000 from 300 - 1,800 m/z. AGC target was set to 4e5 and 50 ms. The top 20 precursors per cycle were subjected to HCD fragmentation at 15,000 resolution with an isolation width of 0.7 m/z, 30 NCE, 3e4 AGC target and 50ms max injection time. For TMT experiments, resolution was set to 60,000 and 34 NCE. Dynamic exclusion was enabled with a duration of 45 sec.

Spectra were searched using Spectrum Mill against the same database as the one used for the MHC I IP/MS spectra analysis (described above), specifying Trypsin/allow P (allows K-P and R-P cleavage) as digestion enzyme, allowing 4 missed cleavages. Carbamidomethylation of cysteine was set as a fixed modification. For the GBM dataset TMT labeling was required at lysine, but peptide N-termini were allowed to be either labeled or unlabeled. Allowed variable modifications were acetylation at the protein N-terminus, oxidized methionine, pyroglutamic acid, deamidated asparagine and pyrocarbamidomethyl cysteine. Match tolerances were set to 20 ppm on MS1 and MS2 level. PSMs score thresholding used the Spectrum Mill auto-validation module to apply target-decoy based FDR in 2 steps: at the peptide spectrum match (PSM) level and the protein level. In step 1 PSM-level autovalidation was done first using an auto-thresholds strategy with a minimum sequence length of 8; automatic variable range precursor mass filtering; and score and delta Rank1 – Rank2 score thresholds optimized to yield a PSM-level FDR estimate for precursor charges 2 through 4 of <1.0% for each precursor charge state in each LC-MS/MS run. To achieve reasonable statistics for precursor charges 5-6, thresholds were optimized to yield a PSM-level FDR estimate of <0.5% across all LC runs per experiment (instead of per each run), since many fewer spectra are generated for the higher charge states. In step 2, protein-polishing autovalidation was applied to each experiment to further filter the PSMs using a target protein-level FDR threshold of zero, the protein grouping method expand subgroups, top uses shared (SGT) with an absolute minimum protein score of 9. After assembling protein groups from the autovalidated PSMs, protein polishing determined the maximum protein level score of a protein subgroup that consisted entirely of distinct peptides estimated to be false-positive identifications (PSMs with negative delta forward-reverse scores); B721: 11.6, GBM: 10.5. PSMs were removed from the set obtained in the initial peptide-level autovalidation step if they contributed to protein subgroups that had protein scores below the maximum false-positive protein score. In Spectrum Mill the protein score was the sum of the scores of distinct peptides. When a peptide sequence of >8 residues was shared by multiple protein entries in the sequence database, the proteins were grouped together. In some cases there were unshared peptides that uniquely represent a subgroup, i.e. lower scoring member of the group, typically isoforms, family members, or different species. As a consequence of these two peptide and protein level steps, each identified protein subgroup was comprised of multiple peptides, unless a single excellent scoring peptide was the sole match.

In the cases where a spectrum could be matched to multiple peptide sequences from different ORFs, the same decision tree was followed for the whole proteome analysis as for the MHC I described above.

### Estimation of absolute translation levels

Our improved translation quantification based on Ribo-seq reads incorporates multi-mapping information and translated frame information. To account for multi-mapping, reads were scaled based on their number of alignments: For example, if a read maps to a 5 different ORFs, it will contribute 0.2 at each location. Using the offset-corrected SAM file generated by RibORF (described above), and given that we know the translated frame identified by Ribo-seq, we counted the total number of multimapping-adjusted reads that are in-frame for each ORF in nuORFdb’s BED12 file using a custom script, and calculated TPM using those read counts and the ORF length. The Python script is provided.

### Peptide sequence correlation, clustering and visualization

Peptide distance computation and visualization were performed as before (Abelin et al. 2017). Briefly, peptide distances were defined as: where *A* is the allele; %_&_and %_’_are peptide sequences; < is the length of the peptide sequences, *n*∈{8,9,10,11}; *H* is the entropy of the amino acid residues at each position in the peptide, -.%/01234 = >?@01234 − 01234 is a 20×20 matrix of residue dissimilarities derived from a pre-computed matrix of residue similarities biased by their HLA binding properties (Y. Kim et al. 2009). For each allele, peptide distances between every pair of peptides in the MS datasets was computed and the pairwise distance matrices were reduced to two dimensions with non-metric multidimensional scaling (NMDS) (nmds() function from ecodist R package).

Peptide sequence motif correlation (for Fig. 2f) was calculated per allele using all detected 9AA peptides. For each peptide, the frequency of each amino acid at each position was calculated to generate a vector of 180 features long. Using these vectors, the position entropy weighted correlation was found between nuORF peptides and all annotated peptides, or between 10,000 random subsets of annotated peptides the same size as the nuORF set and all annotated peptides (minus the subset). Correlations were calculated for all 92 measured alleles independently

### MHC I binding affinity prediction

Fig.2g: HLAthena (http://hlathena.tools/) (Sarkizova et al. 2019) was used to predict MHC I binding affinities for the predicted spliced peptides from Faridi et al (Faridi et al. 2018).

Fig.4d,e: NetMHCpan v4.0 (Jurtz et al. 2017) was used to predict MHC I binding affinities for the HLA alleles expressed in MEL11, to remain consistent with previous studies (Ott et al. 2017).

### Whole genome sequencing and analysis

PCR-free Whole Genome Sequencing (WGS) was performed on cultured melanoma patient 11 cells and matched healthy PBMCs at the Broad Genomics Platform. Libraries were prepared using the Kapa Biosciences HyperPrep library construction kit, and sequenced to 60x coverage (Illumina 2×150bp reads, NovaSeq). Cancer-specific variants were identified using GATK Best Practices (GATK v3.x nightly-2017-09-30) (Bateman et al. 2002) and Strelka2 v2.8.4 (S. Kim et al. 2018). In particular, we first aligned sequenced reads to human genome reference assembly GRCh37 using BWA-MEM (H. Li 2013) v0.7.15-r1140 with default parameters. We then sort aligned reads by coordinates and removed PCR duplicates using Picard tool v2.12.1 [http://broadinstitute.github.io/picard/]. Next, we applied base quality score recalibration to the de-duplicated BAM files using GATK. The recalibrated BAM files were used as inputs for both GATK and Strelka2 for calling somatic variants. For GATK, we followed best practices and used MuTect2 with --dbsnp set to dbSNP build 138 [https://www.ncbi.nlm.nih.gov/snp/] and –cosmic set to Cosmic v82 [https://cosmic-blog.sanger.ac.uk/cosmic-release-v82/]. For Strelka2, we first ran Manta (Chen et al. 2016) v1.2.1 to detect structural variants and indels as recommended by Strelka2 user guide [https://github.com/Illumina/strelka/blob/v2.9.x/docs/userGuide/README.md]. We then ran Strelka2 with --indelCandidates option set to Manta outputs and other options set to default values. We merged variants called using GATK and Strelka2 together.

### Variant analysis, read coverage, and neoantigen predictions

To derive ORFs containing cancer-specific variants identified by WGS, variants that were found within the reference transcripts used in the study were selected using bedtools intersect (Quinlan and Hall 2010) v2.25.0 of the BED12 file of transcripts with the VCF file of variants. Variants were then incorporated into the transcript sequences, and ORFs were re-derived based on the predicted start codon in nuORFdb and the first in-frame stop codon.

To determine Ribo-seq read coverage and nucleotide identity at the SNV sites, pysam pileup (v0.14.1) was used. To obtain read coverage of indels, bowtie (1.2.2) -m 1 -v 0 was used to align raw sequencing reads (after adapter trimming) to a custom FASTA reference that included matched wild-type and indel-containing regions. No multi-mapping reads or mismatches were allowed, such that only variant- or wild-type supporting reads were retained.

Variants supported by at least 9 Ribo-seq reads and >15% of total reads at the locus were used for neoantigen predictions. To obtain potential neoantigens from the mutated variants, all possible 9- and 10-amino acid long peptides were derived from wild-type and variant-containing proteins in nuORFdb. Peptides unique to the variant-containing proteins were retained as potential neoantigens. NetMHCpan v4.0 was used to predict neoantigen binding affinities to HLA alleles (Jurtz et al. 2017). Indels were visualized in IGV (Robinson et al. 2011) to identify in-frame Ribo-seq reads supporting the translation of indel-generated frame-shifted ORFs and wild-type ORFs.

### Identification of tissue-specific or tissue-enriched nuORFs

For the TCGA analysis, we included 473 available skin cutaneous melanoma (SKCM) samples and 172 glioblastoma multiforme (GBM) samples. For GTEx (GTEx Consortium et al. 2017), we randomly selected 10 samples from each tissue. For CLL, available data from 390 CLLs and 21 B-cell samples from healthy donors were included. These comprise two cohorts: 106 CLL and 12 healthy samples from DFCI/Broad Institute (Landau et al. 2015) and 284 CLL and 9 healthy samples from Spanish ICGC studies (Ferreira et al. 2014; Puente et al. 2015). FASTQ files from all cohorts were aligned using STAR v2.6.1d (Dobin et al. 2013) to the reference human genome GRCh37, using the transcriptome annotation combing GENCODE and MiTranscriptome, as used for Ribo-seq based ORF detection described above. Expression at the gene-level was quantified using RNA-SeQC v2.3.3, and expression at the isoform level was quantified using RSEM v1.3.1 (B. Li and Dewey 2011). The parameters used for all components of this pipeline are described at https://github.com/broadinstitute/gtex-pipeline/blob/v9/TOPMed_RNAseq_pipeline.md.

### Identifying cancer-enriched nuORFs based on MHC I IP LC-MS/MS

We generated a list of 335 nuORFs detected by LC-MS/MS in the MHC I immunopeptidomes of the 4 melanoma samples we analyzed. We rank ordered nuORFs by mean expression of the parent transcript across all GTEx samples, excluding the testis, and selected 34 nuORFs with mean expression in the lowest 15% enriching for those not expressed or lowly expressed in healthy tissues. We further filtered them based on the nuORF parent transcript expression across 473 melanoma samples in the TCGA, retaining 6 nuORFs where at least 5% of TCGA samples had expression 2-fold or greater than the highest level detected in any GTEx sample.

### Identifying cancer-specific nuORFs based on Ribo-seq

Based on the Ribo-seq translation levels (TPM) (available through NCBI GEO:GSE143263), we selected nuORFs with TPM > 0 across all in-group samples (all CLL samples / all GBM samples/ all MEL samples) and TPM = 0 in the rest of the Ribo-seq samples profiled. We retained those nuORFs with parent transcript TPM < 1 across healthy tissues in GTEx, excluding the testis.

## Statistical analyses

Fig.2a,c: In the comparison of the MS/MS spectrum scores calculated by Spectrum Mill (Fig. 2a) as well as the translation levels of ORFs (Fig. 2c), the sample sizes were very large, thus the t-tests showed significance, yet the effect size is small, as shown by the confidence intervals calculated using linear regression by the python package statsmodels.regression.linear_model.OLS.

Fig.2d, S***u***pplementary ***Fig.4d: Retention time vs. predicted hydrophobicity:*** Lowess was fit to the annotated peptide retention time and hydrophobicity values using the python package sm.nonparametric.lowess. Residuals between annotated peptide identifications to the lowess fit and residuals between nuORF peptide identifications to the lowess fit were computed and compared with rank sum test in python using scipy.stats.ranksums.

Fig.2h: The lengths of detected Canonical ORFs were compared to the lengths of the detected ORFs in each of the shown categories using a t-test with unequal variance in python using scipy.stats.ttest_ind.

Fig.2m, S***u***pplementary ***Fig.7h:*** The cumulative distribution functions (CDFs) for length or translation level (TPM) of annotated ORFs or nuORFs detected in the MHC-I immunopeptidome or in the whole proteome, compared with a KS test using the python scipy.stats.kstest.

Fig.3g: Given the variable number of known and B721 matched HLA alleles in cancer patients, we simulated the % overlap with variable numbers of alleles matching. All overlaps were measured between 6 B721 alleles randomly sampled from the measured 92 alleles, with a fixed number of type matched alleles. These simulations were calculated for both annotated and nuORF peptides. We then calculated a linear regression between the number of matched alleles and the median % overlap for each cancer sample for both annotated and nuORF.

Fig.4e: Using netMHCpan v4.0, we predicted the rate of strong binders (predicted binding <500 nM) for all high confidence SNVs that also showed strong Ribo-seq support, with at least 9 Ribo-seq reads and 15% of all reads supporting the SNV. We compared the strong binder rate for annotated- and nuORF-derived mutations using a t-test and calculated confidence intervals using linear regression.

Fig.4i: For each nuORF identified as being cancer type specific using ribosome profiling data and low GTEx expression, we compared the expression in TCGA for the associated cancer type to other cancer types and to GTEx, with a rank sum test in python scipy.stats.ranksums. Higher expression in respective TCGA samples was indicated on the far right of 4I and the percent of predicted nuORFs significantly upregulated is shown in 4J.

Fig 2k, S***u***pplementary ***Fig.7c,f:*** We tested for enrichment or depletion of nuORF types in Whole Proteome or cancer samples by generating a % detected distribution for each nuORF type by randomly sampling 1 to 6 B721 alleles from the 92 measured, and reporting the % of nuORFs of each type. We then calculated the p-value for enrichment or depletion as the ratio of the simulated distribution greater than or less than the observed, respectively. To test for overall enrichment or depletion in cancers, we used a t-test to compare the observed p-values to a normal distribution.

## Data and Code availability

Python scripts and Jupyter notebooks used in the analysis are available on GitHub: https://github.com/klarman-cell-observatory/Riboseq-nuORFs.

## Sequencing data

The raw Ribo-seq data (fastq.gz), offset-corrected BAM files used for translated ORF identification by RibORF and BigWig file generation, BigWig files for Ribo-seq data visualization in genome browsers, and Ribo-seq translation levels (TPM) are deposited to NCBI GEO (GSE143263) for established cell lines (B721.221, A375 and HCT116), and for primary melanocytes (Thermo C0025C). GTEx, TCGA, CLL and healthy B cell samples RNA-seq transcription quantification of transcript isoforms is deposited to NCBI GEO: GSE143263. Ribo-seq translation levels (TPM) of primary GBM and melanoma samples are deposited to NCBI GEO: GSE143263. Raw data pertaining to primary patient samples is deposited to dbGaP. B721.221 RNA seq data for HLA-C (C*04:01, C*07:01) is deposited under GEO: GSE131267. Melanoma RNA-seq data are deposited in dbGaP (https://www.ncbi.nlm.nih.gov/projects/gap/cgi-bin/study.cgi?study_id=phs001451.v1.p1 (Ott et al. 2017)). Glioblastoma bulk RNA-seq data are available through dbGaP (https://www.ncbi.nlm.nih.gov/gap) with accession number phs001519.v1.p1 (Keskin et al. 2019).

## Mass spectrometry data

The original mass spectra for immunopeptidomes of 2 melanoma patient-derived cell lines and the full proteome of a glioblastoma patient-derived cell line, tables of peptide spectrum matches for all experiments, and the protein sequence databases used for searches have been deposited in the public proteomics repository MassIVE (https://massive.ucsd.edu) These datasets will be made public upon acceptance of the manuscript. Original mass spectrometry data for the previously published mono-allelic immunopeptidomes, B721.221 cell line full proteome, and patient-derived cell line immunopeptidomes are accessible at ftp://massive.ucsd.edu/MSV000080527, ftp://massive.ucsd.edu/MSV000084172, and ftp://massive.ucsd.edu/MSV000084442. All other data are available from the corresponding authors upon reasonable request.

---

**Sup. Fig. 1.**
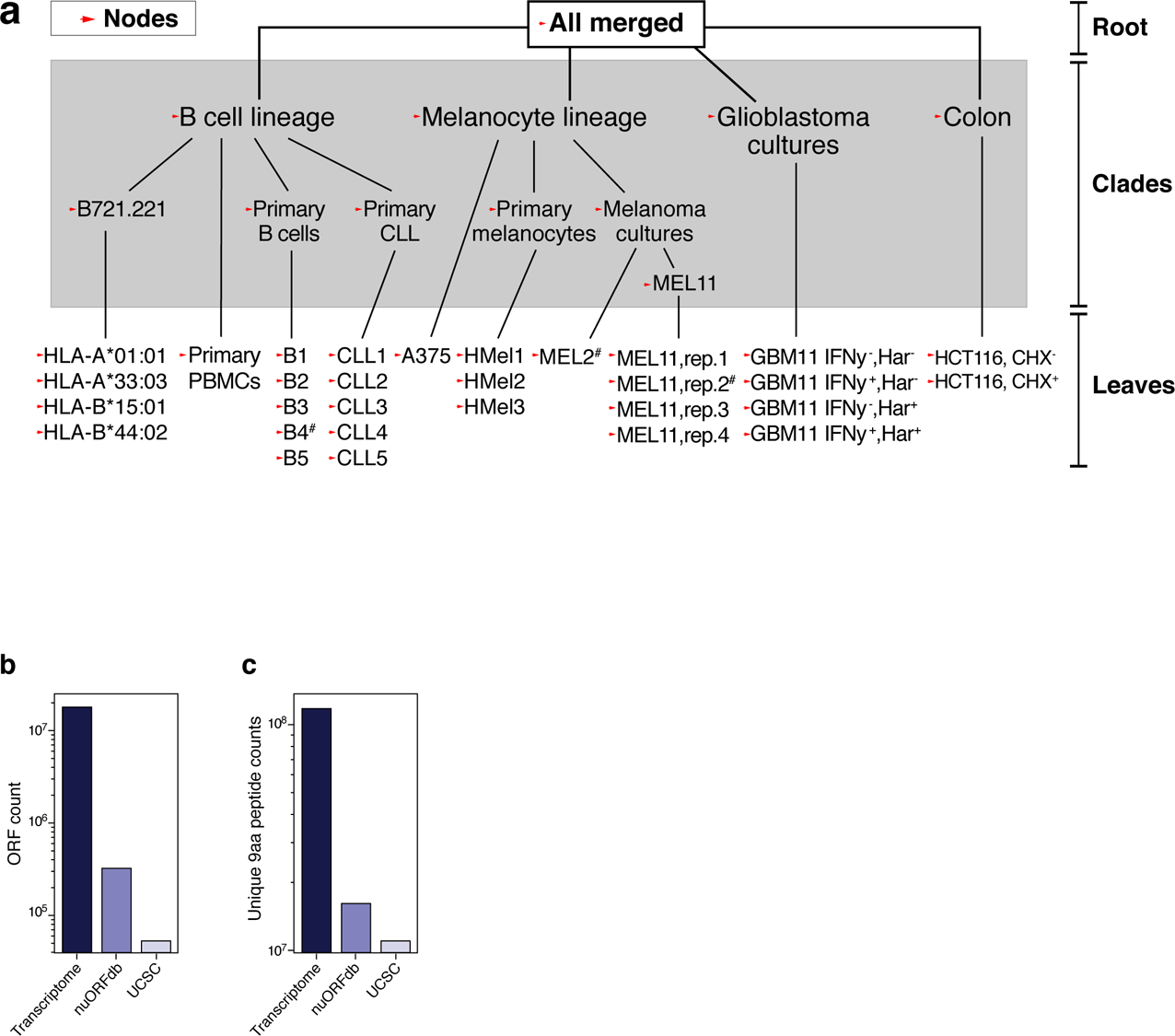
nuORFdb characteristics. **a.** Hierarchical ORF prediction. Tree showing individual samples (leaves), combinations of samples (clades) and entire datasets of all reads (root) representing the nodes used to make ORF predictions (arrowheads). #: samples used in nuORFdb construction, but later discovered to be of poor quality and not used in any subsequent analyses; CHX: samples pre-treated with cycloheximide; Harr: samples pretreated with harringtonine, IFNy: samples pre-treated with interferon gamma. **b,c.** NuORFdb size relative to the transcriptome and the annotated proteome. Number of ORFs (y axis, **b**) and unique 9AA peptides (y axis, **c**) in the entire transcriptome, the nuORFdb, or the annotated UCSC proteome (x axis).

**Sup. Fig. 2.**
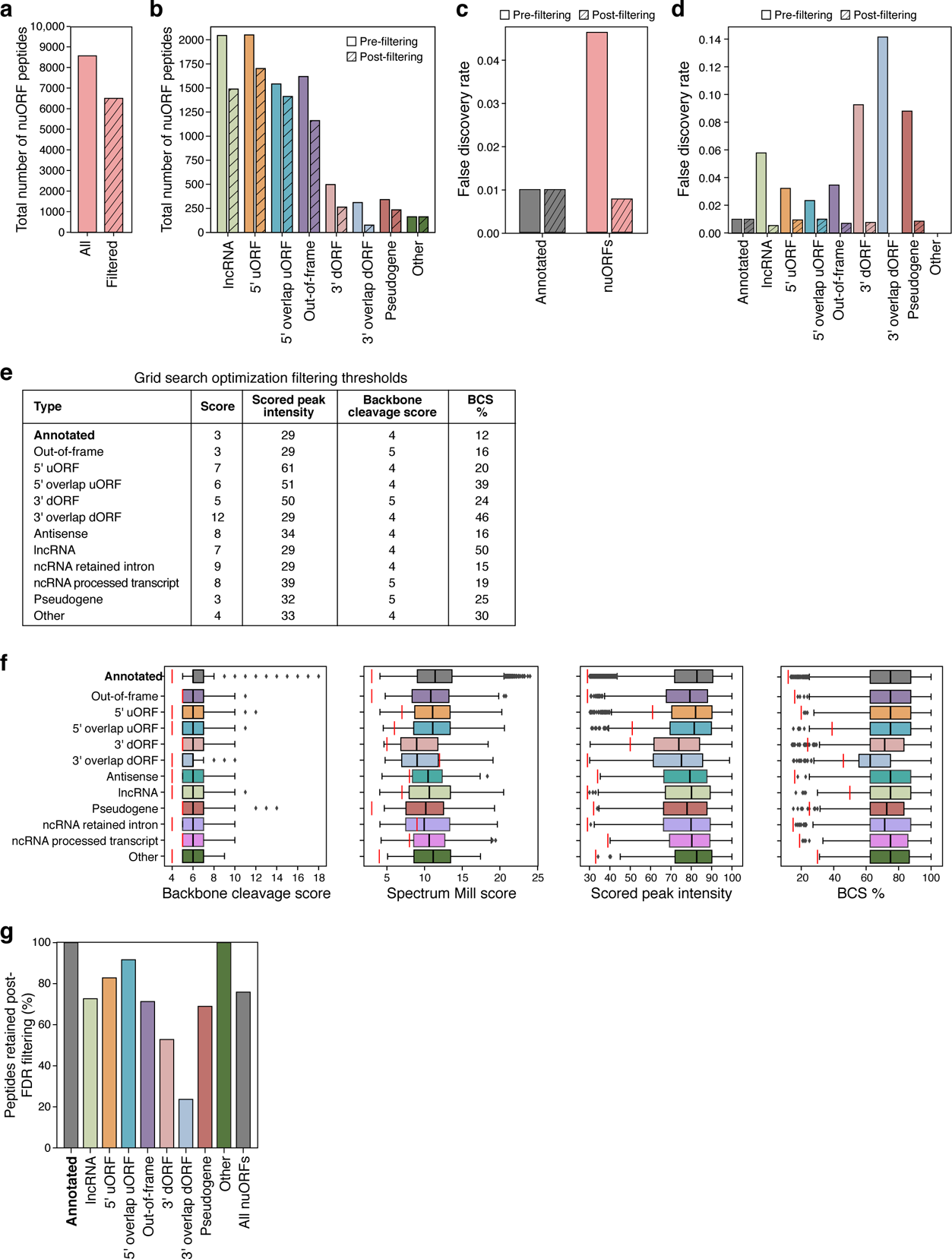
Additional filtering of MHC I IP, MS/MS-detected nuORF peptides. **a-d.** Impact of filtering on nuORF number, types and false discovery rates. **a,b.** Total number of nuORF peptides (y axis) identified pre-filtering (solid bars) and retained post-filtering (hashed bars) overall (**a**) and for different nuORF types (x axis, **b**). **c,d.** False discovery rate (y axis) for annotated (grey) and nuORF (pink) peptides across 92 HLA alleles pre- and post-filtering (hashed) overall (**c**) and for different ORF types (x axis, **d**). **e.** Criteria used to filter peptides across ORF types. **f.** Filtering thresholds across nuORF categories. Filter cutoffs (vertical red lines) across different peptide spectral match scoring features (x axis) for different ORF types (y axis). Median, with 25% and 75% (box range), and 1.5 IQR (whiskers) are shown. **g.** Filtering impact across categories. Percent of peptides (y axis) retained post-filtering across different ORF categories and overall (x axis).

**Sup. Fig. 3.**
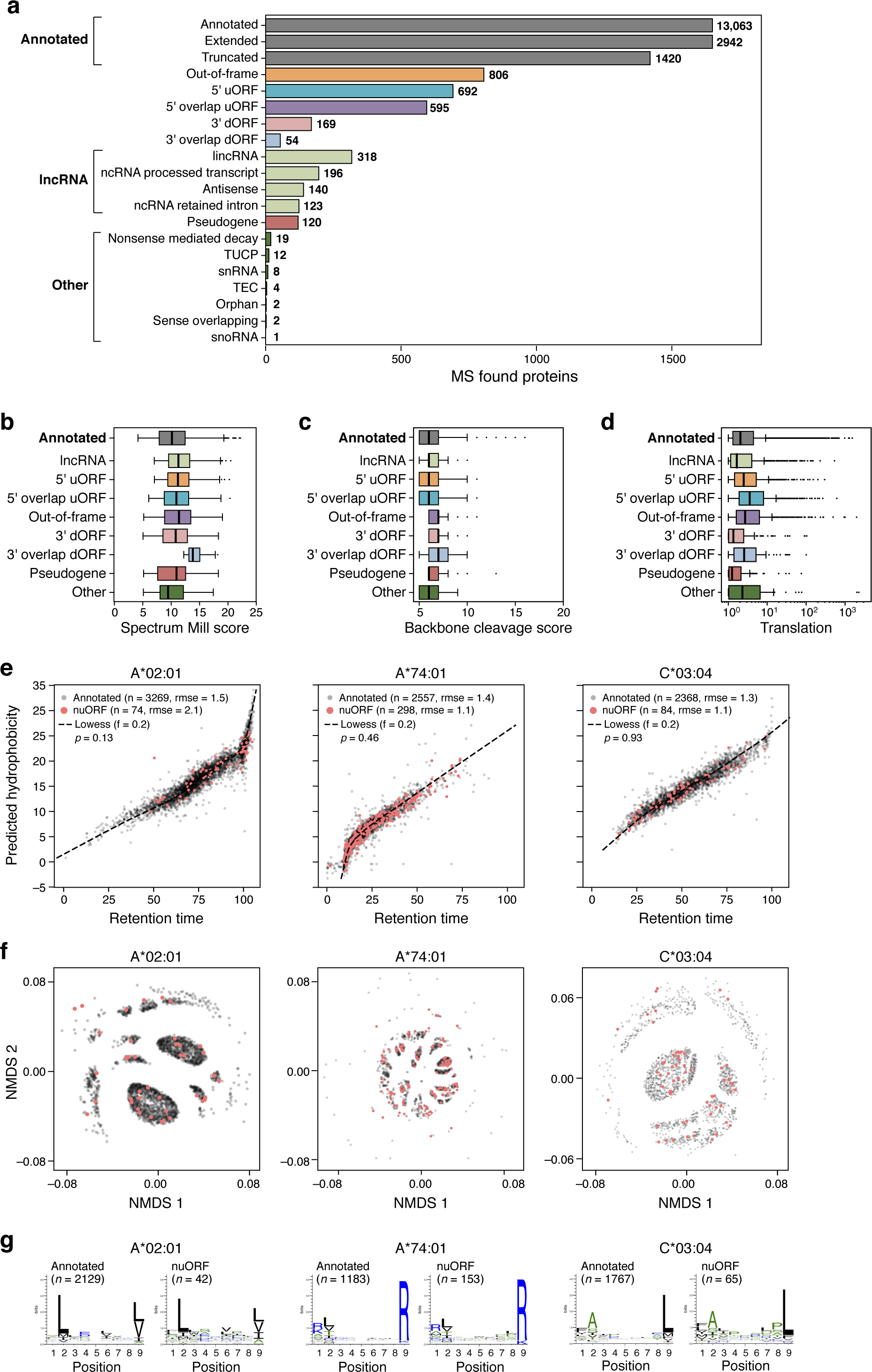
nuORFs peptides in the MHC I immunopeptidome have comparable biochemical properties to annotated peptides. **a.** MHC I immunopeptidome includes peptides from different nuORF categories. Number of unique proteins (x axis) detected by MHC I IP LC-MS/MS across expanded ORF types (y axis). **b-g.** Comparable biochemical features of nuORF and annotated peptides. **b.** Distribution of LC-MS/MS Spectrum Mill identification score (x axis) for annotated and nuORF peptides across ORF types (y axis). **c.** Peptide fragmentation score (x axis) for peptides identified across ORF types (y axis). **d.** Ribo-seq translation levels (x axis, log2(TPM+1)) of MHC I MS-detected ORFs across various ORF types (y axis). For all boxplots, median, with 25% and 75% (box range), and 1.5 IQR (whiskers) are shown. **e.** Predicted hydrophobicity index (y axis) against the LC-MS/MS retention time (x axis) for annotated (grey) and nuORF (pink) peptide sequences for three representative HLA alleles. Dashed line: Lowess fit to the annotated peptides. Sample sizes, root mean square errors (rmse), and p-values (rank-sum test on residuals) are marked. f,g. Similar sequence motifs in nuORFs and annotated peptides. **f.** Non-metric multidimensional scaling (NMDS) plot of all MHC IP LC-MS/MS-detected annotated and nuORF 9 AA peptide sequences clustered by peptide sequence similarity for three representative HLA alleles. **g.** Consensus peptide sequence motif plots of all MHC IP LC-MS/MS-detected annotated and nuORF 9 AA peptide sequences.

**Sup. Fig. 4.**
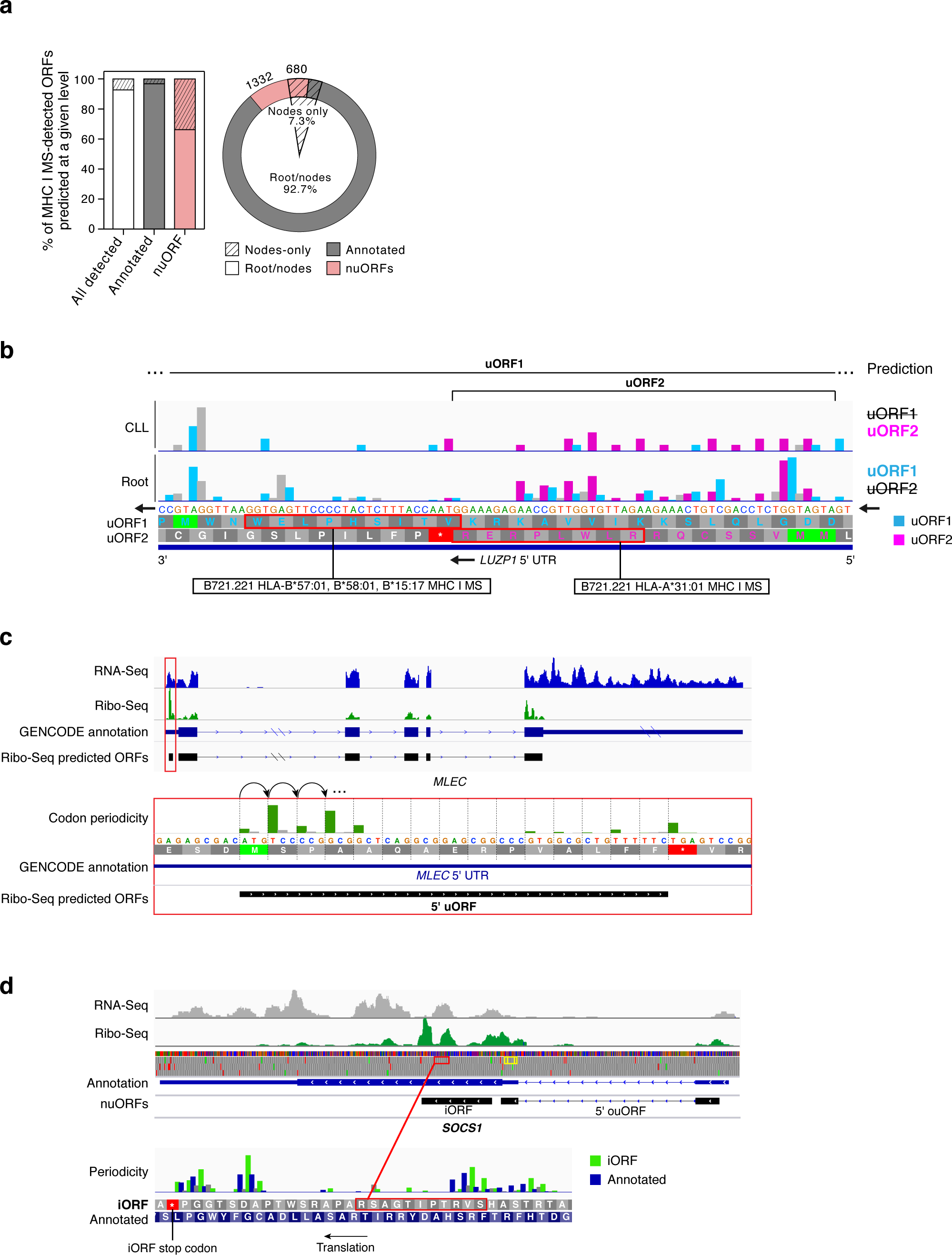
Hierarchical ORF prediction based on Ribo-seq identifies short, overlapping, tissue-specific nuORFs. **a.** nuORFs predictions are more sample and tissue specific than annotated ORFs. Proportion of annotated ORFs (grey) and nuORFs (pink) in the MHC I immunopeptidome (y axis, and pie chart). Hashed: proportion predicted only at the leaf and clade level, but not at the root. **b.** Two overlapping, MHC I MS-detected 5’ uORFs in *LUZP1* as an example of tissue-specific, overlapping nuORFs identified by hierarchical ORF prediction. uORF2 (pink) was predicted in the CLL clade, and not at the root. uORF1 (cyan) was predicted at the root and not in the CLL clade. Detected peptides outlined in red with the HLA alleles where peptides were detected marked below. **c.** Example of identification of short ORFs proximal to long annotated ORFs in the *MLEC* gene. RNA-seq (blue) and Ribo-seq (green) reads aligned to the transcript of the *MLEC* gene. RNA-seq reads align to the entire length of the transcript, while Ribo-seq reads align exclusively to the translated portions. Ribo-seq supports translation of a 5’ uORF (red box, top). Histogram of reads supporting translation of the *MLEC* 5’ uORF (dark green) (bottom). **d.** *SOCS1* gene as an example of identification of short, overlapping nuORFs. *SOCS1* gene encodes three translated proteins: the annotated ORF, an out-of-frame iORF, and a 5’ overlap ouORF. Two MHC I MS-detected peptides from 5’ ouORF outlined in yellow. Detected iORF peptide outlined in red and shown in higher magnification below. Bottom: Histogram of Ribo-seq reads supporting translation of the annotated ORF (blue) and the out-of-frame iORF (green).

**Sup. Fig. 5.**
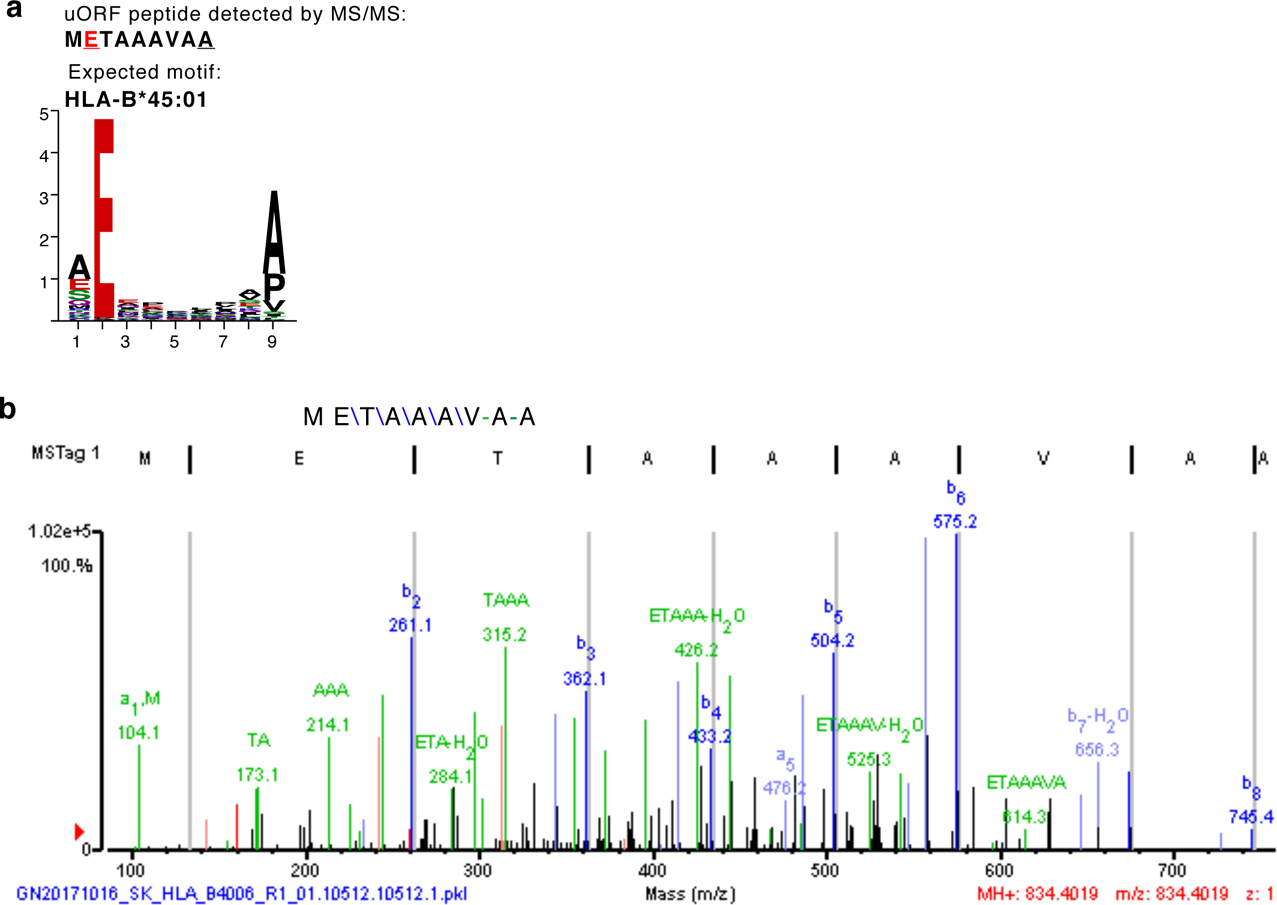
Short nuORFs are presented on MHC I without post-translational protease processing. **a.** The peptide comprising the full-length sequence of the 5’ uORF from the *ARAF* gene and the expected binding motif for the HLA allele B*45:01, from which it was presented and detected. **b.** LC-MS/MS spectrum of the *ARAF* 5’ uORF.

**Sup. Fig. 6.**
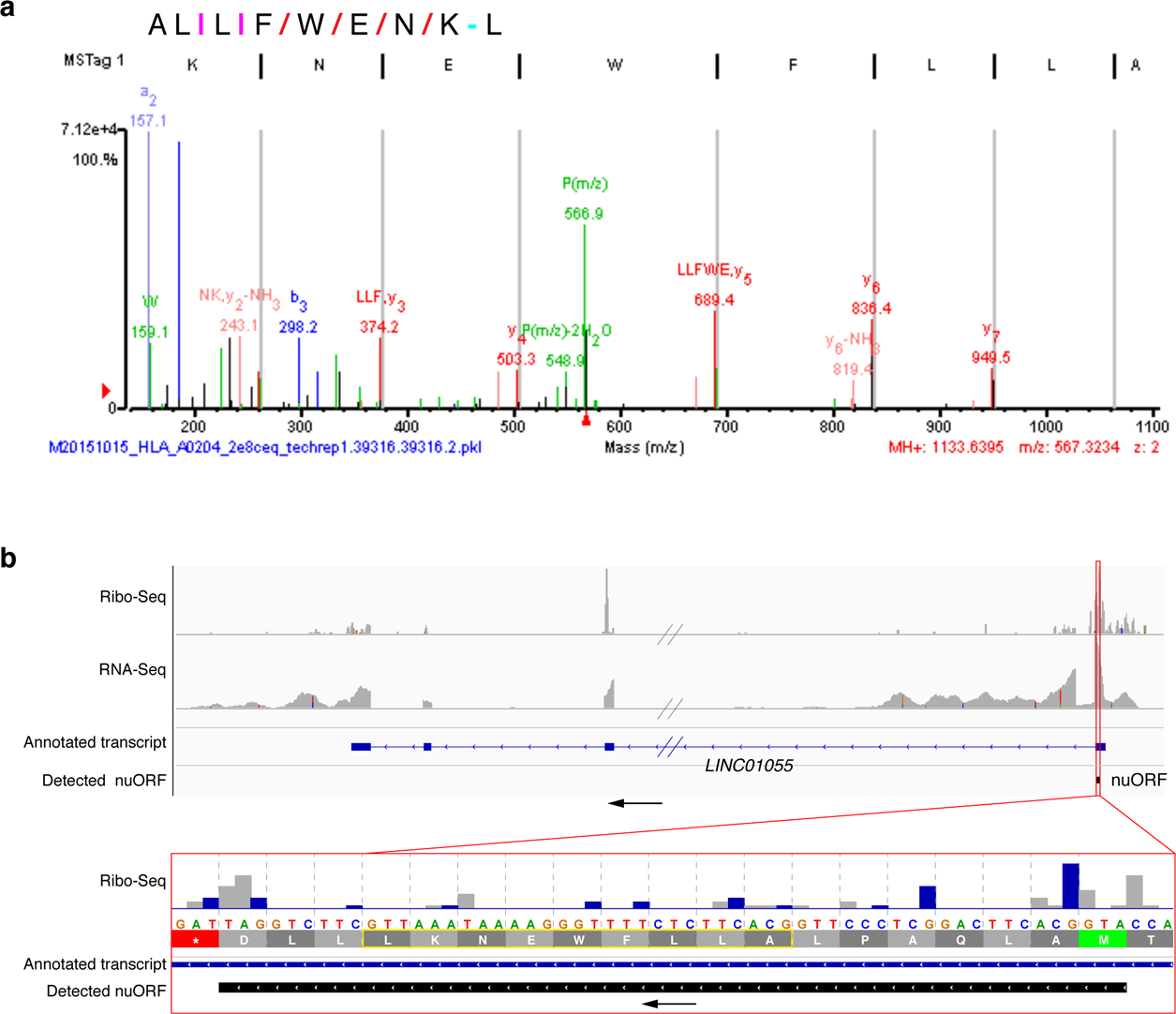
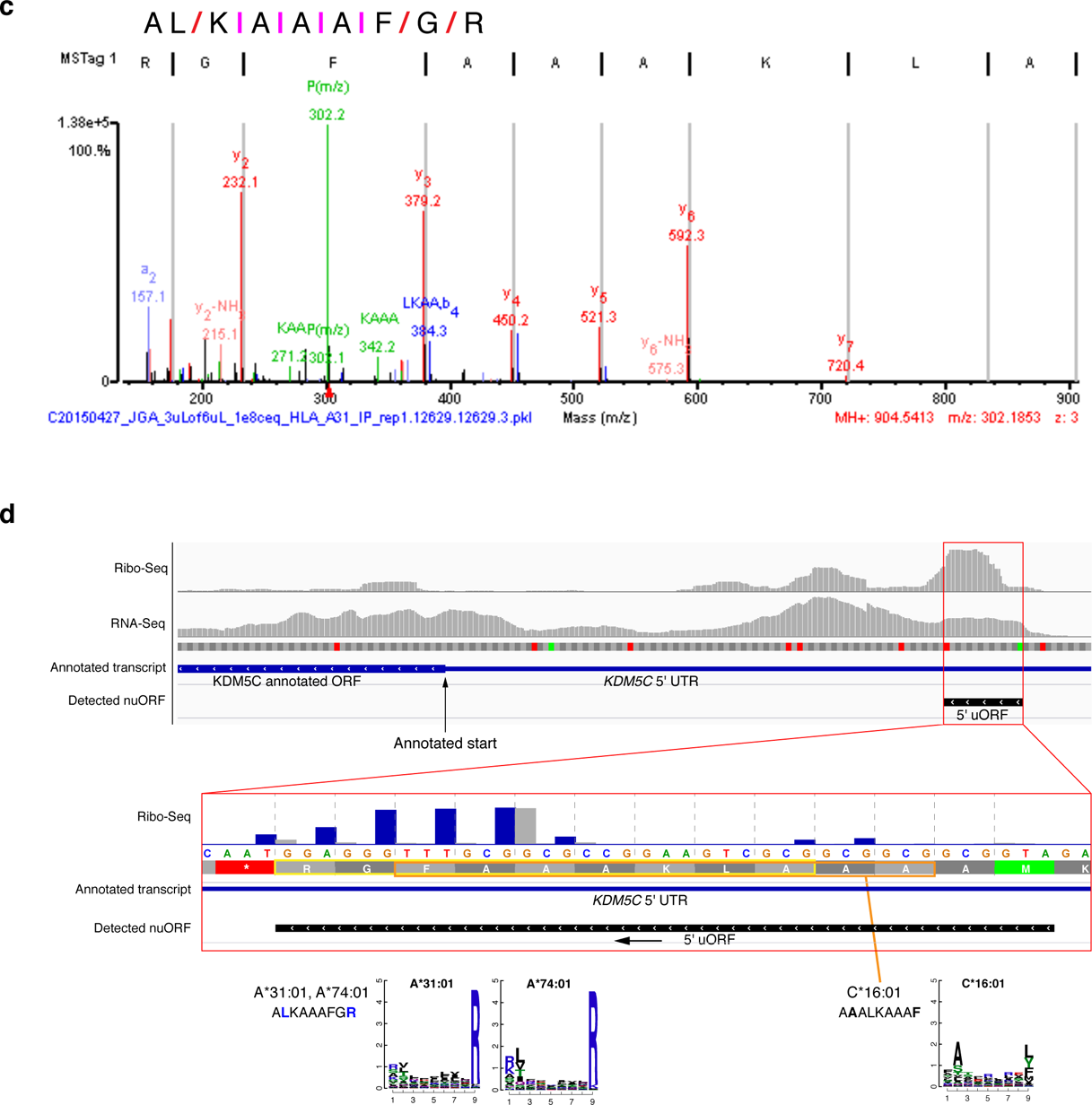
Some nuORF-derived peptides map to the same MS/MS spectra as peptides that have previously been proposed to be derived from proteasomal splicing in Faridi et al. **a.** MS/MS spectrum of the peptide ALLFWENKL presented by HLA allele A*02:04 that can be derived from the translated *LINC01055* lncRNA nuORF was previously proposed to be derived from proteasomal cis-splicing. The peptide contains L at both position 2 and the C-terminus, consistent with the anchor motif for allele A*02:04 ligands. **b.** RNA-seq and Ribo-seq reads aligned to the *LINC01055* lncRNA locus. Red box marks the MHC I IP LC-MS/MS detected nuORF. Bottom panel shows a magnified view of the reads supporting nuORF translation. Detected peptide is outlined in yellow. **c,d.** Partial sequence present in the MS/MS spectra assigned to spliced peptides are also consistent with different, yet similar, nuORF peptide sequences. **c.** MS/MS spectrum of a peptide presented by allele A*31:01 yields near complete fragmentation with explicit sequence evidence for the order of all residues except the first two: AL/K|A|A|A|F/G/R. Both peptides ALKAAAFGR (derived from *KDM5C* 5’ uORF) and LAKAAAFGR (proposed to be derived from proteasomal cis-splicing) are consistent with the fragment ions present in the spectrum. Leucine in position 2 (nuORF) and arginine at the C-terminus are consistent with the anchor motif for allele A*31:01 ligands, while alanine in position 2 (spliced) is not. In addition, 3 more peptides from *KDM5C* 5’ uORF were detected on HLA A*74:01 and C*16:01, further supporting the nuORF translation and presentation. **d.** RNA-seq and Ribo-seq reads aligned to the *KDM5C* locus. Red box marks the 5’ uORF detected by MHC I IP LC-MS/MS. Detected peptides and the expected peptide sequence motifs are outlined in yellow and orange.

**Sup. Fig. 7.**
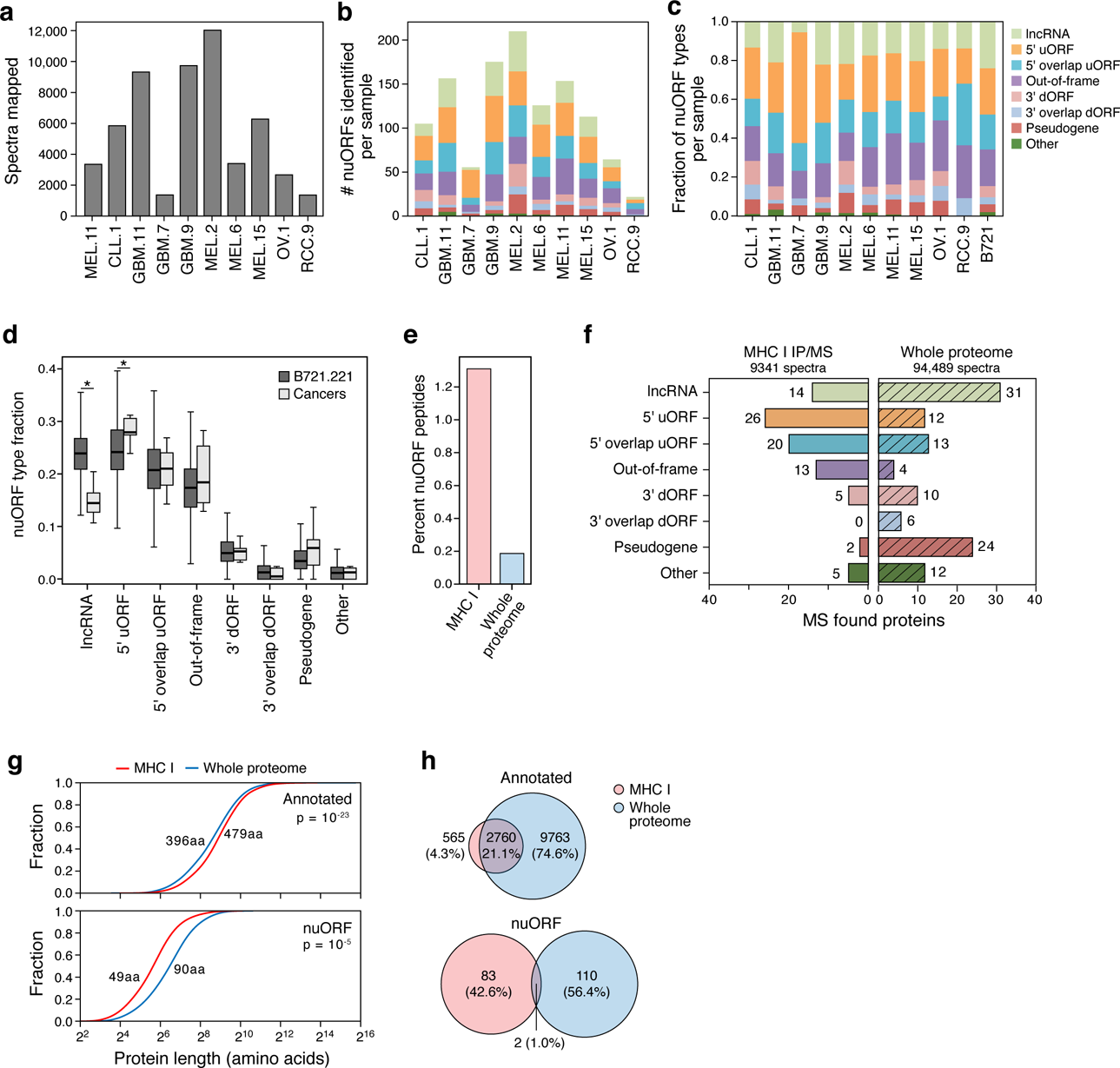
nuORF peptides in the MHC I immunopeptidome and whole proteome of cancer cells. **a.** nuORFdb helps map immunopeptidome even from samples and tumor types not used in constructing the reference. Total number of MHC I LC-MS/MS spectra mapped (y axis) across cancer samples (x axis). **b-d.** nuORFs of various types were detected in the MHC I immunopeptidome of cancer samples. Number (**b**) and proportion (**c**) of nuORFs (y axis) of different types identified in each cancer sample (x axis). **d.** Distribution of the fraction (y axis) of nuORF types (x axis) in B721.221 cells (dark grey) or across cancer samples (light grey). Asterisk: p < 0.05, rank-sum test. Median, with 25% and 75% (box range), and 1.5 IQR (whiskers) are shown. **e-h.** nuORFs are more abundant in the MHC I immunopeptidome than in the whole proteome. **e.** Percent of nuORF peptides (y axis) detected in the immunopeptidome (pink) and in the whole proteome (blue) of GBM11. **f.** Number of nuORFs (x axis) of different types (y axis) identified in the MHC I immunopeptidome (left) vs. whole proteome (hatched, right) in GBM11. **g.** Protein length (x axis, amino acids) of annotated (top) and nuORF (bottom) proteins detected in the MHC I immunopeptidome (pink) vs. in the whole proteome (blue). p-values: KS test. **h.** Proportion of all annotated ORFs (top) or nuORFs (bottom) detected in the whole proteome (blue), immunopeptidome (pink) or both (intersection) in GBM11.

**Sup. Fig. 8.**
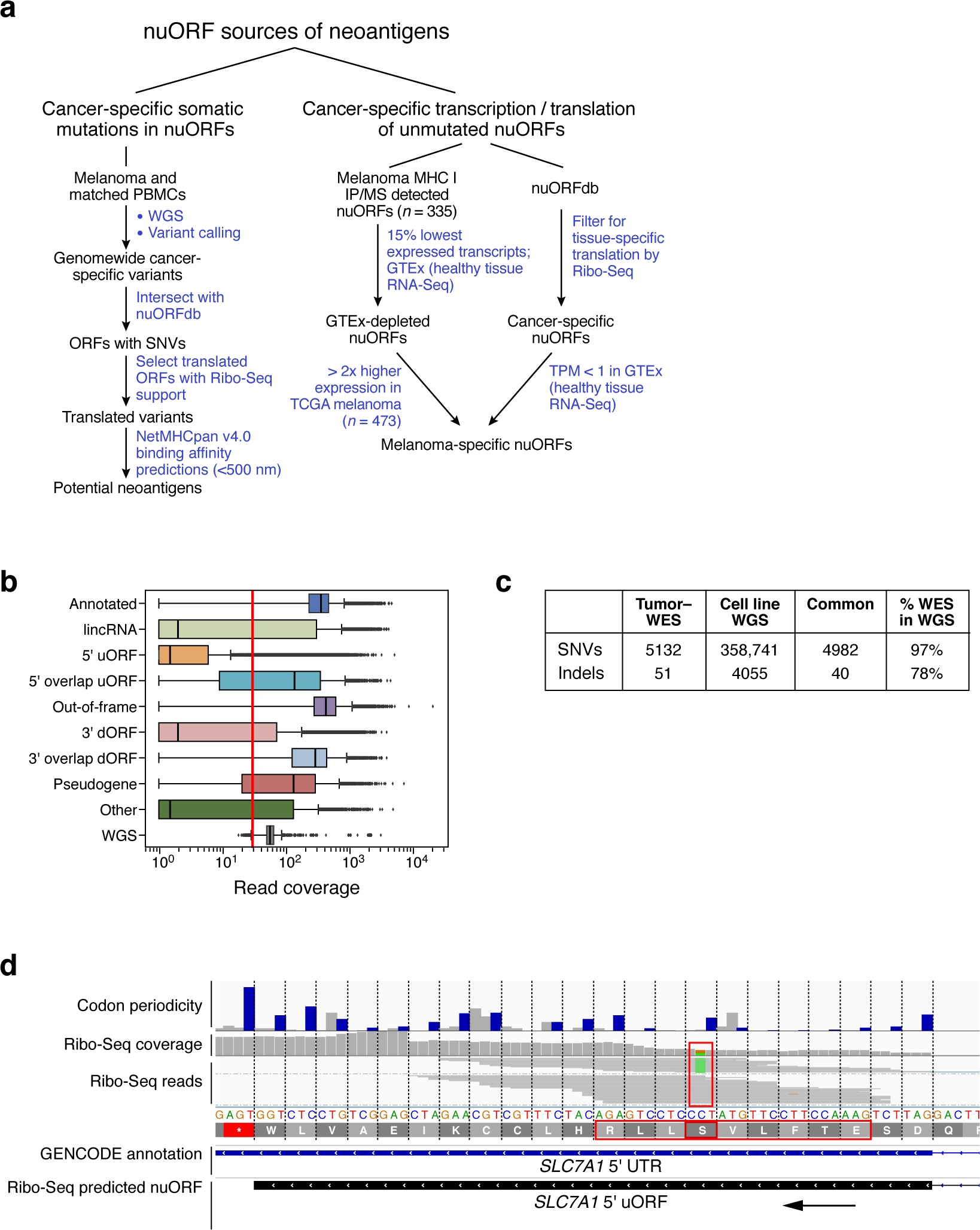
nuORFs can be potential sources of neoantigens. **a.** Approaches to identify potential nuORF-derived neoantigens. **b.** nuORFs have low sequence coverage by WES compared to WGS. Distribution of WES read coverage (x axis) across different ORF types (y axis). Bottom: WGS read coverage across all ORFs of all types. Vertical red line marks 30× coverage. Median, with 25% and 75% (box range), and 1.5 IQR (whiskers) are shown. **c.** Somatic variants in the melanoma patient-derived cell line reflect the variants detected in the original tumor. Cancer-specific SNVs and InDels identified by WES from the primary tumor and by WGS from the tumor-derived cell line. **d.** Ribo-seq can be used to identify translated variants. Example of a translated *SLC7A1* 5’ uORF with a cancer-specific SNV. Top: histogram of Ribo-seq reads supporting the translation of the 5’ uORF. Middle: Ribo-seq reads supporting translation of the mutant (green) and wild-type alleles. Predicted neoantigen outlined in red.

**Sup. Fig. 9.**
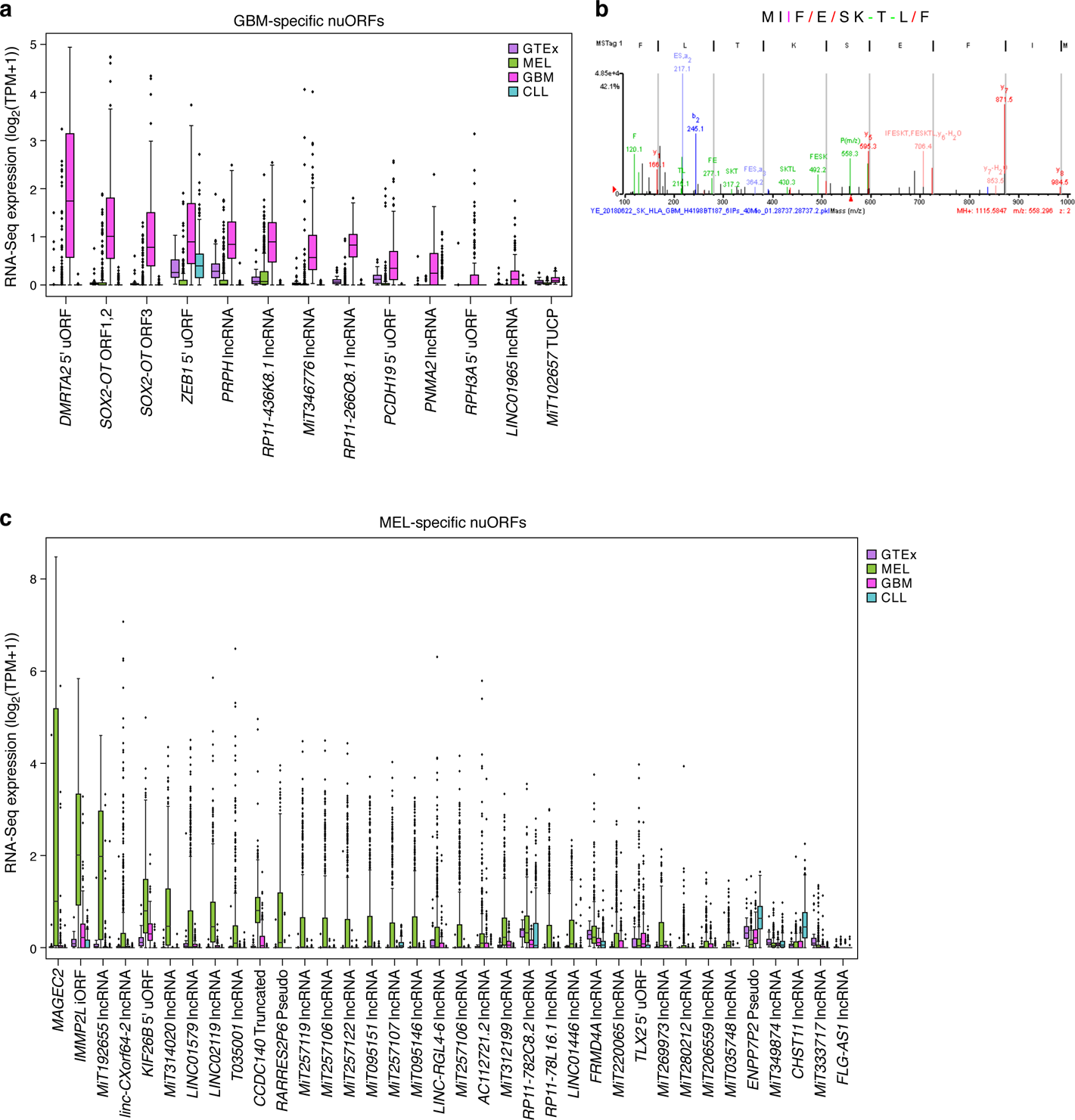
GBM and melanoma specific nuORFs. **a-b.** GBM-specific nuORFs. **a.** RNA-seq expression (y axis, log2(TP-M+1)) of GBM-specific nuORFs (x axis) in GTEx and tumor samples. **b.** LC-MS/MS spectrum of a peptide from *SOX2-OT* nuORF. **c.** Melanoma-specific nuORFs. RNA-seq expression (y axis, log2(TPM+1)) of melanoma-specific nuORFs (x axis) in GTEx and tumor samples. For all boxplots, median, with 25% and 75% (box range), and 1.5 IQR (whiskers) are shown.

## References

Abelin, J. G., D. B. Keskin, S. Sarkizova, C. R. Hartigan, W. Zhang, J. Sidney, J. Stevens, et al. 2017. “Mass Spectrometry Profiling of HLA-Associated Peptidomes in Mono-Allelic Cells Enables More Accurate Epitope Prediction.” Immunity 46 (2): 315–26.

Blum, Amy, Peggy Wang, and Jean C. Zenklusen. 2018. “SnapShot: TCGA-Analyzed Tumors.” Cell 173 (2): 530.

Chew, Guo-Liang, Andrea Pauli, John L. Rinn, Aviv Regev, Alexander F. Schier, and Eivind Valen. 2013. “Ribosome Profiling Reveals Resemblance between Long Non-Coding RNAs and 5’ Leaders of Coding RNAs.” Development 140 (13): 2828–34.

Consortium, G. TEx. 2015. “Human Genomics. The Genotype-Tissue Expression (GTEx) Pilot Analysis: Multitissue Gene Regulation in Humans.” Science 348 (6235): 648–60.

Erhard, F., A. Halenius, C. Zimmermann, A. L’Hernault, D. J. Kowalewski, M. P. Weekes, S. Stevanovic, R. Zimmer, and L. Dolken. 2018. “Improved Ribo-Seq Enables Identification of Cryptic Translation Events.” Nature Methods. https://doi.org/10.1038/nmeth.4631.

Faridi, Pouya, Chen Li, Sri H. Ramarathinam, Julian P. Vivian, Patricia T. Illing, Nicole A. Mifsud, Rochelle Ayala, et al. 2018. “A Subset of HLA-I Peptides Are Not Genomically Templated: Evidence for Cis- and Trans-Spliced Peptide Ligands.” Science Immunology 3 (28). https://doi.org/10.1126/sciimmunol.aar3947.

Fields, A. P., E. H. Rodriguez, M. Jovanovic, N. Stern-Ginossar, B. J. Haas, P. Mertins, R. Raychowdhury, et al. 2015. “A Regression-Based Analysis of Ribosome-Profiling Data Reveals a Conserved Complexity to Mammalian Translation.” Molecular Cell 60 (5): 816–27.

Frankish, A., M. Diekhans, A. M. Ferreira, R. Johnson, I. Jungreis, J. Loveland, J. M. Mudge, et al. 2019. “GENCODE Reference Annotation for the Human and Mouse Genomes.” Nucleic Acids Research 47 (D1): D766–73.

Georgiadis, Panagiotis, Irene Liampa, Dennie G. Hebels, Julian Krauskopf, Aristotelis Chatziioannou, Ioannis Valavanis, Theo M. C. M. de Kok, et al. 2017. “Evolving DNA Methylation and Gene Expression Markers of B-Cell Chronic Lymphocytic Leukemia Are Present in Pre-Diagnostic Blood Samples More than 10 Years prior to Diagnosis.” BMC Genomics 18 (1): 728.

Hilf, Norbert, Sabrina Kuttruff-Coqui, Katrin Frenzel, Valesca Bukur, Stefan Stevanović, Cécile Gouttefangeas, Michael Platten, et al. 2019. “Actively Personalized Vaccination Trial for Newly Diagnosed Glioblastoma.” Nature 565 (7738): 240–45.

Hoof, Ilka, Bjoern Peters, John Sidney, Lasse Eggers Pedersen, Alessandro Sette, Ole Lund, Søren Buus, and Morten Nielsen. 2009. “NetMHCpan, a Method for MHC Class I Binding Prediction beyond Humans.” Immunogenetics 61 (1): 1–13.

Hutter, Carolyn, and Jean Claude Zenklusen. 2018. “The Cancer Genome Atlas: Creating Lasting Value beyond Its Data.” Cell 173 (2): 283–85.

Hu, Zhuting, Patrick A. Ott, and Catherine J. Wu. 2018. “Towards Personalized, Tumour-Specific, Therapeutic Vaccines for Cancer.” Nature Reviews. Immunology 18 (3): 168–82.

Ingolia, N. T., S. Ghaemmaghami, J. R. Newman, and J. S. Weissman. 2009. “Genome-Wide Analysis in Vivo of Translation with Nucleotide Resolution Using Ribosome Profiling.” Science 324 (5924): 218–23.

Iyer, M. K., Y. S. Niknafs, R. Malik, U. Singhal, A. Sahu, Y. Hosono, T. R. Barrette, et al. 2015. “The Landscape of Long Noncoding RNAs in the Human Transcriptome.” Nature Genetics 47 (3): 199– 208.

Ji, Z., R. Song, A. Regev, and K. Struhl. 2015. “Many lncRNAs, 5’UTRs, and Pseudogenes Are Translated and Some Are Likely to Express Functional Proteins.” eLife 4. https://doi.org/10.7554/eLife.08890.

Keskin, D. B., A. J. Anandappa, J. Sun, I. Tirosh, N. D. Mathewson, S. Li, G. Oliveira, et al. 2019. “Neoantigen Vaccine Generates Intratumoral T Cell Responses in Phase Ib Glioblastoma Trial.” Nature 565 (7738): 234–39.

Laumont, C. M., T. Daouda, J. P. Laverdure, E. Bonneil, O. Caron-Lizotte, M. P. Hardy, D. P. Granados, et al. 2016. “Global Proteogenomic Analysis of Human MHC Class I-Associated Peptides Derived from Non-Canonical Reading Frames.” Nature Communications 7: 10238.

Laumont, C. M., K. Vincent, L. Hesnard, E. Audemard, E. Bonneil, J. P. Laverdure, P. Gendron, et al. 2018. “Noncoding Regions Are the Main Source of Targetable Tumor-Specific Antigens.” Science Translational Medicine 10 (470). https://doi.org/10.1126/scitranslmed.aau5516.

Liepe, Juliane, Fabio Marino, John Sidney, Anita Jeko, Daniel E. Bunting, Alessandro Sette, Peter M. Kloetzel, Michael P. H. Stumpf, Albert J. R. Heck, and Michele Mishto. 2016. “A Large Fraction of HLA Class I Ligands Are Proteasome-Generated Spliced Peptides.” Science 354 (6310): 354–58.

Ma, J., J. K. Diedrich, I. Jungreis, C. Donaldson, J. Vaughan, M. Kellis, J.R. Yates3rd, and A. Saghatelian. 2016. “Improved Identification and Analysis of Small Open Reading Frame Encoded Polypeptides.” Analytical Chemistry 88 (7): 3967–75.

Ma, J., C. C. Ward, I. Jungreis, S. A. Slavoff, A. G. Schwaid, J. Neveu, B. A. Budnik, M. Kellis, and A. Saghatelian. 2014. “Discovery of Human sORF-Encoded Polypeptides (SEPs) in Cell Lines and Tissue.” Journal of Proteome Research 13 (3): 1757–65.

Martinez, Thomas F., Qian Chu, Cynthia Donaldson, Dan Tan, Maxim N. Shokhirev, and Alan Saghatelian. 2019. “Accurate Annotation of Human Protein-Coding Small Open Reading Frames.” Nature Chemical Biology, December. https://doi.org/10.1038/s41589-019-0425-0.

Mylonas, Roman, Ilan Beer, Christian Iseli, Chloe Chong, Hui-Song Pak, David Gfeller, George Coukos, Ioannis Xenarios, Markus Müller, and Michal Bassani-Sternberg. 2018. “Estimating the Contribution of Proteasomal Spliced Peptides to the HLA-I Ligandome.” Molecular & Cellular Proteomics: MCP 17 (12): 2347–57.

Ott, P. A., Z. Hu, D. B. Keskin, S. A. Shukla, J. Sun, D. J. Bozym, W. Zhang, et al. 2017. “An Immunogenic Personal Neoantigen Vaccine for Patients with Melanoma.” Nature. https://doi.org/10.1038/nature22991.

Rajasagi, M., S. A. Shukla, E. F. Fritsch, D. B. Keskin, D. DeLuca, E. Carmona, W. Zhang, et al. 2014. “Systematic Identification of Personal Tumor-Specific Neoantigens in Chronic Lymphocytic Leukemia.” Blood 124 (3): 453–62.

Raj, A., S. H. Wang, H. Shim, A. Harpak, Y. I. Li, B. Engelmann, M. Stephens, Y. Gilad, and J. K. Pritchard. 2016. “Thousands of Novel Translated Open Reading Frames in Humans Inferred by Ribosome Footprint Profiling.” eLife 5. https://doi.org/10.7554/eLife.13328.

Robbins, P. F., M. El-Gamil, Y. F. Li, E. B. Fitzgerald, Y. Kawakami, and S. A. Rosenberg. 1997. “The Intronic Region of an Incompletely Spliced gp100 Gene Transcript Encodes an Epitope Recognized by Melanoma-Reactive Tumor-Infiltrating Lymphocytes.” Journal of Immunology 159 (1): 303–8.

Rodríguez, Ana Eugenia, Jose Ángel Hernández, Rocío Benito, Norma C. Gutiérrez, Juan Luis García, María Hernández-Sánchez, Alberto Risueño, et al. 2012. “Molecular Characterization of Chronic Lymphocytic Leukemia Patients with a High Number of Losses in 13q14.” PloS One 7 (11): e48485.

Rolfs, Zach, Markus Müller, Michael R. Shortreed, Lloyd M. Smith, and Michal Bassani-Sternberg. 2019. “Comment on ‘A Subset of HLA-I Peptides Are Not Genomically Templated: Evidence for Cis- and Trans-Spliced Peptide Ligands.’” Science Immunology 4 (38). https://doi.org/10.1126/sciimmunol.aaw1622.

Sahin, U., E. Derhovanessian, M. Miller, B. P. Kloke, P. Simon, M. Lower, V. Bukur, et al. 2017. “Personalized RNA Mutanome Vaccines Mobilize Poly-Specific Therapeutic Immunity against Cancer.” Nature 547 (7662): 222–26.

Sarkizova, Siranush, Susan Klaeger, Phuong M. Le, Letitia W. Li, Giacomo Oliveira, Hasmik Keshishian, Christina R. Hartigan, et al. 2020. “A Large Peptidome Dataset Improves HLA Class I Epitope Prediction across Most of the Human Population.” Nature Biotechnology, 38, 199–209. https://doi.org/10.1038/s41587-019-0322-9.

Su, Rui, Shuo Cao, Jun Ma, Yunhui Liu, Xiaobai Liu, Jian Zheng, Jiajia Chen, et al. 2017. “Knockdown of SOX2OT Inhibits the Malignant Biological Behaviors of Glioblastoma Stem Cells via up-Regulating the Expression of miR-194-5p and miR-122.” Molecular Cancer 16 (1): 171.

Van Den Eynde, B. J., B. Gaugler, M. Probst-Kepper, L. Michaux, O. Devuyst, F. Lorge, P. Weynants, and T. Boon. 1999. “A New Antigen Recognized by Cytolytic T Lymphocytes on a Human Kidney Tumor Results from Reverse Strand Transcription.” The Journal of Experimental Medicine 190 (12): 1793–1800.

Wang, R. F., S. L. Johnston, G. Zeng, S. L. Topalian, D. J. Schwartzentruber, and S. A. Rosenberg. 1998. “A Breast and Melanoma-Shared Tumor Antigen: T Cell Responses to Antigenic Peptides Translated from Different Open Reading Frames.” Journal of Immunology 161 (7): 3596–3606.

Yewdell, J. W. 2011. “DRiPs Solidify: Progress in Understanding Endogenous MHC Class I Antigen Processing.” Trends in Immunology 32 (11): 548–58.

Yoshimura, Akihiko, Tetsuji Naka, and Masato Kubo. 2007. “SOCS Proteins, Cytokine Signalling and Immune Regulation.” Nature Reviews. Immunology 7 (6): 454–65.

## References

Bassani-Sternberg, Michal, Eva Bräunlein, Richard Klar, Thomas Engleitner, Pavel Sinitcyn, Stefan Audehm, Melanie Straub, et al. 2016. “Direct Identification of Clinically Relevant Neoepitopes Presented on Native Human Melanoma Tissue by Mass Spectrometry.” Nature Communications 7 (November): 13404.

Bateman, Alex, William R. Pearson, Lincoln D. Stein, Gary D. Stormo, and John R. Yates III, eds. 2002. “From FastQ Data to High-Confidence Variant Calls: The Genome Analysis Toolkit Best Practices Pipeline.” In Current Protocols in Bioinformatics, 467:11.10.1–11.10.33. Hoboken, NJ, USA: John Wiley & Sons, Inc.

Chen, Xiaoyu, Ole Schulz-Trieglaff, Richard Shaw, Bret Barnes, Felix Schlesinger, Morten Källberg, Anthony J. Cox, Semyon Kruglyak, and Christopher T. Saunders. 2016. “Manta: Rapid Detection of Structural Variants and Indels for Germline and Cancer Sequencing Applications.” Bioinformatics 32 (8): 1220–22.

Dobin, Alexander, Carrie A. Davis, Felix Schlesinger, Jorg Drenkow, Chris Zaleski, Sonali Jha, Philippe Batut, Mark Chaisson, and Thomas R. Gingeras. 2013. “STAR: Ultrafast Universal RNA-Seq Aligner.” Bioinformatics 29 (1): 15–21.

Ferreira, Pedro G., Pedro Jares, Daniel Rico, Gonzalo Gómez-López, Alejandra Martínez-Trillos, Neus Villamor, Simone Ecker, et al. 2014. “Transcriptome Characterization by RNA Sequencing Identifies a Major Molecular and Clinical Subdivision in Chronic Lymphocytic Leukemia.” Genome Research 24 (2): 212–26.

GTEx Consortium, Laboratory, Data Analysis &Coordinating Center (LDACC)—Analysis Working Group, Statistical Methods groups—Analysis Working Group, Enhancing GTEx (eGTEx) groups, NIH Common Fund, NIH/NCI, NIH/NHGRI, et al. 2017. “Genetic Effects on Gene Expression across Human Tissues.” Nature 550 (7675): 204–13.

Jurtz, Vanessa, Sinu Paul, Massimo Andreatta, Paolo Marcatili, Bjoern Peters, and Morten Nielsen. 2017. “NetMHCpan-4.0: Improved Peptide-MHC Class I Interaction Predictions Integrating Eluted Ligand and Peptide Binding Affinity Data.” Journal of Immunology 199 (9): 3360–68.

Keskin, Derin B., Annabelle J. Anandappa, Jing Sun, Itay Tirosh, Nathan D. Mathewson, Shuqiang Li, Giacomo Oliveira, et al. 2019. “Neoantigen Vaccine Generates Intratumoral T Cell Responses in Phase Ib Glioblastoma Trial.” Nature 565 (7738): 234–39.

Kim, Sangtae, Konrad Scheffler, Aaron L. Halpern, Mitchell A. Bekritsky, Eunho Noh, Morten Källberg, Xiaoyu Chen, et al. 2018. “Strelka2: Fast and Accurate Calling of Germline and Somatic Variants.” Nature Methods 15 (8): 591–94.

Kim, Yohan, John Sidney, Clemencia Pinilla, Alessandro Sette, and Bjoern Peters. 2009. “Derivation of an Amino Acid Similarity Matrix for Peptide: MHC Binding and Its Application as a Bayesian Prior.” BMC Bioinformatics 10 (November): 394.

Krokhin, O. V., Stephen Ying, Rob Craig, Victor Spicer, Werner Ens, K. G. Standing, R. C. Beavis, and J. A. Wilkins. 2004. “B. New Sequence-Specific Correction Factors for Prediction of Peptide Retention in RP--HPLC: Application to Protein Identification by off-Line HPLC-MALDI-MS.” In Proceedings of the 52-Th ASMS Conference on Mass Spectrometry and Allied Topics. Nashville, TN, USA. https://www.researchgate.net/profile/John_Wilkins/publication/265422254_New_sequence-specific_correction_factors_for_prediction_of_peptide_retention_in_RP-HPLC_application_to_protein_identification_by_off-line_HPLC-MALDI-MS/links/5730cf8308aed286ca0dc01b.pdf.

Landau, Dan A., Eugen Tausch, Amaro N. Taylor-Weiner, Chip Stewart, Johannes G. Reiter, Jasmin Bahlo, Sandra Kluth, et al. 2015. “Mutations Driving CLL and Their Evolution in Progression and Relapse.” Nature 526 (7574): 525–30.

Langmead, Ben, Cole Trapnell, Mihai Pop, and Steven L. Salzberg. 2009. “Ultrafast and Memory-Efficient Alignment of Short DNA Sequences to the Human Genome.” Genome Biology 10 (3): R25.

Li, Bo, and Colin N. Dewey. 2011. “RSEM: Accurate Transcript Quantification from RNA-Seq Data with or without a Reference Genome.” BMC Bioinformatics 12 (August): 323.

Li, Heng. 2013. “Aligning Sequence Reads, Clone Sequences and Assembly Contigs with BWA-MEM.” arXiv [q-bio.GN]. arXiv. http://arxiv.org/abs/1303.3997.

Malone, B., I. Atanassov, F. Aeschimann, X. Li, H. Grosshans, and C. Dieterich. 2017. “Bayesian Prediction of RNA Translation from Ribosome Profiling.” Nucleic Acids Research 45 (6): 2960–72.

Mertins, Philipp, Lauren C. Tang, Karsten Krug, David J. Clark, Marina A. Gritsenko, Lijun Chen, Karl R. Clauser, et al. 2018. “Reproducible Workflow for Multiplexed Deep-Scale Proteome and Phosphoproteome Analysis of Tumor Tissues by Liquid Chromatography-Mass Spectrometry.” Nature Protocols 13 (7): 1632–61.

Puente, Xose S., Silvia Beà, Rafael Valdés-Mas, Neus Villamor, Jesús Gutiérrez-Abril, José I. Martín-Subero, Marta Munar, et al. 2015. “Non-Coding Recurrent Mutations in Chronic Lymphocytic Leukaemia.” Nature 526 (7574): 519–24.

Quinlan, Aaron R., and Ira M. Hall. 2010. “BEDTools: A Flexible Suite of Utilities for Comparing Genomic Features.” Bioinformatics 26 (6): 841–42.

Robinson, James T., Helga Thorvaldsdóttir, Wendy Winckler, Mitchell Guttman, Eric S. Lander, Gad Getz, and Jill P. Mesirov. 2011. “Integrative Genomics Viewer.” Nature Biotechnology 29 (1): 24– 26.

Sarkizova, Siranush, Susan Klaeger, Phuong M. Le, Letitia W. Li, Giacomo Oliveira, Hasmik Keshishian, Christina R. Hartigan, et al. 2019. “A Large Peptidome Dataset Improves HLA Class I Epitope Prediction across Most of the Human Population.” Nature Biotechnology, December. https://doi.org/10.1038/s41587-019-0322-9.

